# A dynamic spike threshold with correlated noise predicts observed patterns of negative interval correlations in neuronal spike trains

**DOI:** 10.1101/2020.11.30.403725

**Authors:** Robin S. Sidhu, Erik C. Johnson, Douglas L. Jones, Rama Ratnam

## Abstract

Negative correlations in the sequential evolution of interspike intervals (ISIs) are a signature of memory in neuronal spike-trains. They provide coding benefits including firing-rate stabilization, improved detectability of weak sensory signals, and enhanced transmission of information by improving signal-to-noise ratio. Primary electrosensory afferent spike-trains in weakly electric fish fall into two categories based on the pattern of SCCs: non-bursting units have negative SCCs which remain negative but decay to zero with increasing lags (Type I SCCs), and bursting units have oscillatory (alternating sign) SCCs which damp to zero with increasing lags (Type II SCCs). Here, we predict and match observed ISI serial correlations in these afferents using a stochastic dynamic threshold model. We determine SCCs as a function of an arbitrary discrete noise correlation function **R**_*k*_, where *k* is a multiple of the mean ISI. The function permits forward and inverse calculations of SCCs. Both types of SCCs can be generated by adding colored noise to the spike threshold with Type I SCCs generated with slow noise and Type II SCCs generated with fast noise. We show that a first-order autoregressive (AR) process with a single parameter is sufficient to predict and accurately match both types of afferent SCCs, the type being determined by the sign of the AR parameter. The predicted and experimentally observed SCCs are in geometric progression. The theory predicts that the limiting sum of SCCs is −0.5 yielding a perfect DC-block in the power spectrum of the spike train. Observed SCCs from afferents have a limiting sum that is slightly larger at −0.475 ± 0.04 (mean ± s.d.). We conclude that the underlying process for generating ISIs may be a simple combination of low-order autoregressive (AR) processes, and discuss the results from the perspective of optimal coding.

## 1 Introduction

The spiking activity of many neurons exhibit memory, which stabilizes the neurons’ firing rate and makes it less variable than a renewal process. In spontaneously active neurons, a signature of these memory effects can be found in the serial correlations of interspike intervals (ISIs), which display a prominent negative correlation between adjacent ISIs. This is a result of long intervals following short intervals so that fluctuations from the mean ISI are damped over long time-scales, thereby stabilizing the firing rate [1,2]. Negative correlation between adjacent ISIs, which is the first serial correlation coefficient (*ρ*_1_), can assume a range of values [3] from near-zero (close to a renewal spike train, e.g., [4,5]) to values close to −0.9 [1]. While more negative values may suggest a stronger memory effect, the relationship between the extent of memory in the spike train and their ISI correlations is by no means clear, in part due to the difficulty in determining joint interval distributions of arbitrarily high orders [1,6].

Negative correlations in spontaneous spike trains, or in spike trains obtained under quiescent conditions, have been known for many years. Early reports came from lateral line units of Japanese eel [7], the retina of the cat [8], and subsequently from several neuronal systems [3–5,9–15] (see [3,16] for tabulations of negative correlations). In the active sense of some wave-type weakly electric fish [11,13], primary electrosensory afferents exhibit the strongest known correlations in their adjacent ISIs [1]. These electrosensory neurons are an excellent model system for studying memory effects and regularity in firing due to their high spontaneous spike-rates [1,6,17]. It is likely that spike encoders which demonstrate negative ISI correlations have adaptive value because they can facilitate optimal detection of weak sensory signals [1,2,11,17–21] and enhance information transmission [22–24].

Negative ISI correlations are a characteristic feature of spike frequency adaptation where a constant DC-valued input to a neuron is gradually forgotten, possibly due to an intrinsic high-pass filtering mechanism in the encoder [25–27]. Two commonly used models of spike frequency adaptation are based on: 1) a dynamic threshold, and 2) an adaptation current, both of which cause increased refractoriness following an action potential. In the case of a dynamic threshold, the increased refractoriness is due to an elevation in firing threshold, historically referred to as “accommodation” [28]. This is usually modeled as an abrupt increase in spike threshold immediately after a spike is output, followed by a relaxation of the threshold to its normative value without reset [17,29–37]. Thus, if two spikes are output in quick succession with an interval smaller than the mean ISI, the upward threshold shift will be cumulative, making the membrane more refractory, and so a third spike in the sequence will occur with an ISI that is likely to be longer than the mean. In this way, a dynamic time-varying threshold serves as a moving barrier, carrying with it a memory of prior spiking activity. In the case of an adaptation current, outward potassium currents, including voltage gated potassium currents and calcium-dependent or AHP currents, can give rise to increased refractoriness and lead to spike frequency adaptation (e.g., [5,25–27,36,38,39]). Adaptation currents are usually modeled by explicitly introducing a hyperpolarizing outward current with a time-varying conductance in the equation for the membrane potential. The conductance (or a gating variable) will be elevated following a spike and, as with a dynamic threshold, it will decay back to its normative value. Both models have been successful in reproducing nearly identical spike frequency adaptation behaviour for a step input current, and first serial correlation coefficient (*ρ*_1_), although they differ in other details [27].

In this report we focus on a dynamic threshold model. A simple dynamic threshold model can accurately predict spike-times in cortical neurons [40–42] and peripheral sensory neurons [42]. Further, we had recently proposed that spike trains encoded with a dynamic threshold are optimal, in the sense that they provide an optimal estimate of the input by minimizing coding error (estimation error) for a fixed long-term spike-rate (energy consumption) [42–44]. These results did not incorporate noise in the model, and so here, we extend our earlier model by incorporating noise to model spike timing variability and serial ISI correlations.

In previous work, colored noise or Gaussian noise (or both) are added to a dynamic threshold, or to an adaptation current, or to the input signal so that negative correlations can be observed [17,34,38,45]. In these models the first SCC *ρ*_1_ (between adjacent ISIs) is close to or equal to −0.5, and all remaining correlation coefficients *ρ_i_*, *i* ≥ 2, are identically zero. In another report [27] *ρ*_1_ is parameterized, and can assume values between 0 and −0.5. Experimental spike trains demonstrate broader trends, where *ρ*_1_ can assume values smaller or greater than −0.5, and the remaining SCCs can be non-zero for several lags, sometimes with damped oscillations and sometimes monotonically increasing to zero [1]. Lindner and co-workers investigated several types of integrate-and-fire models with adaptation currents [36,37], adding colored and Gaussian noise to introduce noise fluctuations of different time-scales (slow and fast, respectively). They showed that the time-scale of noise fluctuations determined the various patterns of SCCs, including positive correlations. All of these patterns had a geometric structure (i.e., *ρ_k_*/*ρ*_*k*–1_ = constant). Urdapilleta [46] also obtained a geometric structure with monotonically decaying SCC pattern with *ρ_k_* < 0. These latter studies show that the role of noise fluctuations, in particular the time-scale of fluctuations, are important in determining patterns of SCCs. So, far adaptation models with an adaptation current have successfully predicted patterns of ISIs, but models with dynamic thresholds have been investigated much less, and it is not known whether they can accurately predict observed ISI patterns. In general, models that can accurately predict experimentally observed SCCs for all lags have the potential to isolate mechanisms responsible for ISI fluctuations and negative correlations, and provide insights into neural coding. This is the goal of the current work.

Serial interspike interval correlations observed in primary P-type afferents of weakly electric fish *Apteronotus leptorhynchus* are modeled using a dynamic threshold with noise. The model is analytically tractable and permits an explicit closed-form expression for SCCs in terms of an arbitrary correlation function **R**_*k*_, where *k* is a multiple of the mean ISI. This allows us to solve the inverse problem where we can determine **R**_*k*_ given a sequence of observed SCCs. Theoretically, the limiting sum of the SCCs is −0.5 (a perfect DC-block), and experimental SCCs are close to this sum. This model is parsimonious, and in addition to predicting spike-times as shown earlier, it reproduces observed ISI SCCs. Finally, the model provides a fast method for generating surrogate spike trains that match a mean firing rate with prescribed ISI distribution, joint-ISI distribution, and SCCs.

## 2 Methods

### 2.1 Experimental procedures

Spike data from P-type primary electrosensory afferents were collected in a previous *in vivo* study in *Apteronotus leptorhynchus,* a wave-type weakly electric fish [1]. All animal procedures including animal handling, anesthesia, surgery, and euthanasia received institutional ethics (IACUC) approval from the University of Illinois at Urbana-Champaign, and were carried out as previously reported [1]. No new animal procedures were performed during the course of this work. Of relevance here are some details on the electrophysiological recordings. Briefly, action potentials (spikes) were isolated quasi-intracellularly from P-type afferent nerve fibers in quiescent conditions under ongoing electric organ discharge (EOD) activity (this is the so-called baseline spiking activity of P-type units). An artificial threshold was applied to determine spike onset times, and reported at the ADC sampling rate of 16.67 kHz (sampling period of 60 *μ*s). The fish’s ongoing quasi-sinusoidal EOD was captured whenever possible to determine the EOD frequency from the power spectrum, or the EOD frequency was estimated from the power spectrum of the baseline spike train. Both methods reported almost identical EOD frequencies. EOD frequencies typically ranged from 750 – 1000 Hz (see [1] for more details).

### 2.2 Data analysis

P-type units fire at most once every EOD cycle and this forms a convenient time-base to resample the spike train [1]. Spike times resampled at the EOD rate are reported as increasing integers. Resampling removes the phase jitter in spike-timing but retains long-term correlations due to memory effects in the spike train. Spike-times were converted to a sequence of interspike intervals, *X*_1_ *X*_2_,…, *X_k_*,…, *X_N_*.

#### 2.2.1 Autocorrelation function

The normalized autcorrelation function for the sequence of ISIs are the serial correlation coefficients or SCCs, *ρ*_1_, *ρ*_2_, *ρ*_3_,…, with *ρ*_0_ = 1. These were estimated from time-stamps at the original ADC rate of 16.67 kHz and at the resampled EOD frequency [1]. In some afferents there is a small difference between the two estimates, particularly in the estimates of *ρ*_1_, but this has negligible effect on the results. SCCs were estimated from the resampled spike trains as follows:

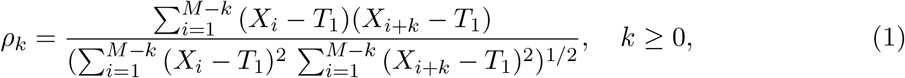

where *T*_1_ is the mean ISI, and *M* is the number of spikes in a block, typically ranging from 1000 – 3000 spikes. The SCCs were estimated in non-overlapping blocks and then averaged over [*N/M*] blocks. There was no drift in the mean ISI within a block. We usually had about 2.5 × 10^5^ spikes per afferent.

#### 2.2.2 Partial autocorrelation function

In addition to the autocorrelation, we compute the partial autocorrelation *ϕ_k,k_* between *X_i_* and *X_i+k_* by removing the linear influence of the intervening variables *X*_*i*+1_,…, *X*_*i*+*k*–1_ [47]. The notation *ϕ_k,j_* means that the process is purely autoregressive (AR) of order *k*, and *ϕ_k,j_* is the *j*^th^ coefficient in the AR model. Partial autocorrelations provide a convenient way to identify an AR process, just as the autocorrelation function, Eq. 1, provides a way to identify a moving average (MA) process. When the ISIs are AR of order-*p*, then the partial correlation function is finite with *ϕ_k,k_* = 0, for *k* > *p*, however, the autcorrelation function will be infinite. Conversely, when the process is MA of order-*m*, then the autocorrelation function is finite with *ρ_k_* = 0, for *k* > *m*, however, the partial autocorrelation function will be infinite. When the partial autocorrelation and autocorrelaton functions are both infinite, then the underlying process is neither purely AR nor purely MA, but is an autoregressive moving average (ARMA) process of some unknown order (*p, m*). The partial autocorrelations *ϕ*_1,1_, *ϕ*_2,2_, *ϕ*_3,3_,…, can be obtained for *k* = 1, 2, 3,…, by solving the Yule-Walker equations. A more efficient method is to solve Durbin’s recursive equations. Durbin’s formula is [47],

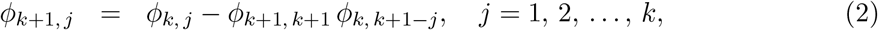

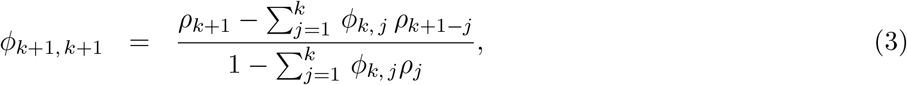

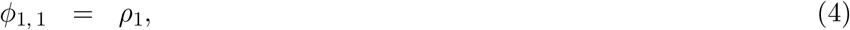

where the *ρ_j_* are SCCs obtained from Eq. 1. The *ϕ_k,j_* in the formula are estimates with some mean and standard deviation over the population. To reduce the estimation error, we can follow the same procedure as for serial correlations by averaging over blocks.

## 3 Results

### 3.1 Experimental results

Serial correlation coefficients (SCCs) were estimated from the baseline activity of 52 P-type primary electrosensory afferents in the weakly electric fish *Apteronotus leptorhynchus* (see Materials and Methods). SCCs from two example afferents (Fig. 1) demonstrate the patterns of negative SCCs observed in spike trains, and motivate this work. Statistical properties of these and other spike trains were reported earlier [1], with a qualitative description of SCCs and some descriptive statistics. A complete analytical treatment is undertaken here. Two types of serial interspike interval statistics can be identified according to the value taken by *ρ*_1_ (the first SCC, between adjacent ISIs).

1. Type I: −0.5 < *ρ*_1_ < 0. Subsequent *ρ_k_* are negative and diminish to 0 with increasing *k* (Fig. 1A). The ISIs of these afferents are unimodal (shown later) and their spike trains do not exhibit bursting activity.
2. Type II: *ρ*_1_ < −0.5. Subsequent *ρ_k_* alternate between positive (*ρ*_2*k*_) and negative (*ρ*_2*k*+1_) values, are progressively damped, and diminish to zero (Fig. 1C). The ISIs of these spike trains are bimodal with a prominent peak at an ISI equal to about one period of the electric organ discharge (EOD) (shown later). These spike trains exhibit strong bursting.

**Figure 1.**
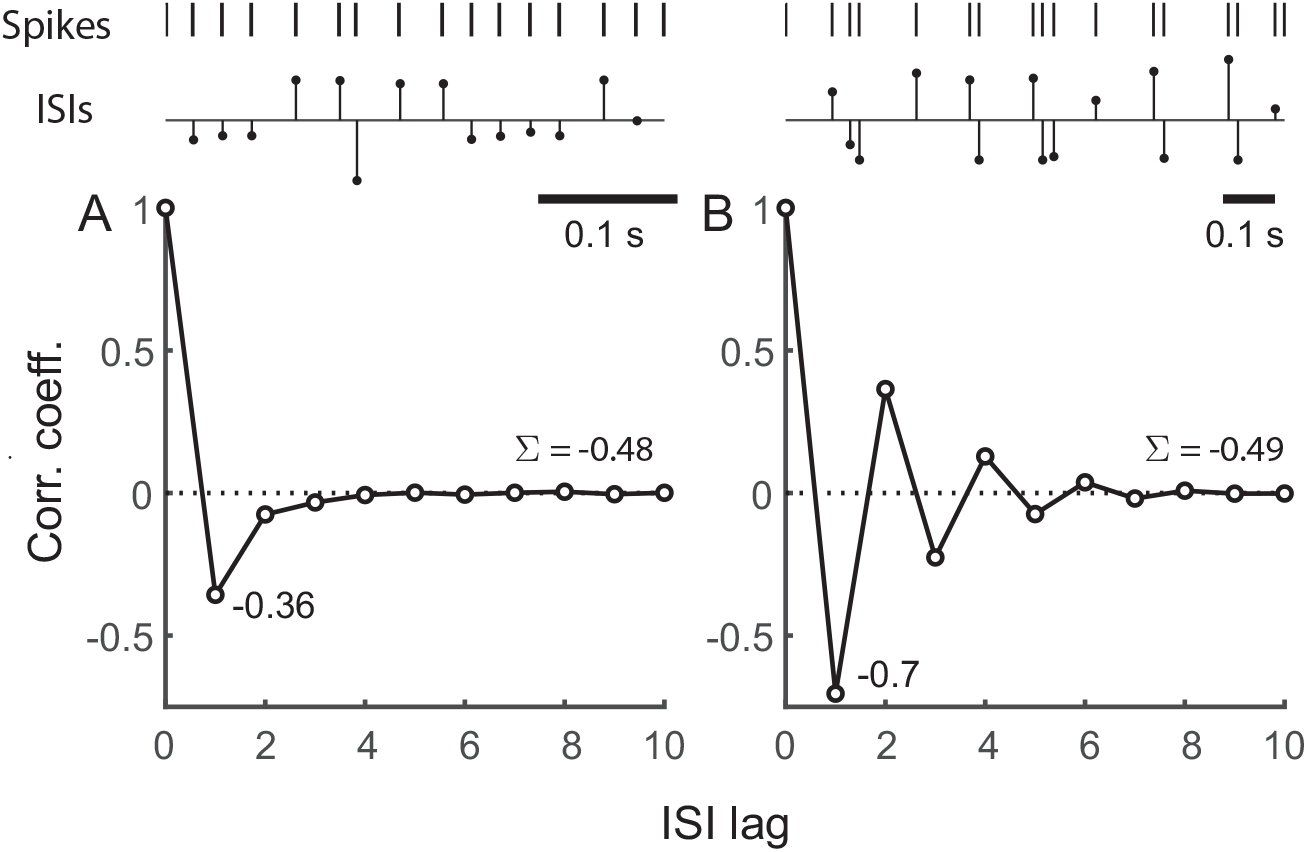
Representative serial interspike interval correlations (SCCs) from two P-type primary electrosensory afferent spike trains from two fish. For each column, from top to bottom, panels depict a sample stretch of spikes, sequence of ISIs for the sample spikes (normalized to mean ISI), and the pattern of ISI correlations. A. Type I (*ρ*_1_ > −0.5): Non-bursting unit with first SCC *ρ*_1_ = −0.36. Remaining *ρ_k_* < 0 diminish to 0. Sum of SCCs (∑) over 15 lags is −0.48. B. Type II (*ρ*_1_ < −0.5): Strongly bursting unit with *ρ*_1_ = −0.7 with marked alternating positive and negative correlations. Sum of SCCs (∑) over 15 lags is −0.49. Spike trains sampled at 60 μs. Mean ISI ± SD (in ms): 2.42 ± 0.72 (A), and 6.04 ± 3.59 (B). Electric organ discharge (EOD) frequency: 948 Hz (A), and 990 Hz (B).

Afferent fibers sampled from individual fish were a mix of Types I and II. Additionally we identify a third type that has not been observed in experimental data (at least by these authors) but is fairly common in some models with adaptation (e.g., [17,34], see also Discussion). We call this Type III, and for this pattern of SCCs *ρ*_1_ = −0.5 and subsequent *ρ_k_* are identically zero (Fig. 1). The Type III pattern is a singleton (i.e., there is only one SCC pattern in this class).

SCCs for observed Type I and Type II units *ρ_k_* (*k* ≥ 1) sum to approximately −0.5. For Type III, the sum is exactly −0.5.

The baseline spike-train statistics of 52 P-type afferents reported earlier [1] were analyzed in detail in this work. All observed units showed a negative correlation between succeeding ISIs (*ρ*_1_ < 0) (Fig. 2A). Experimental spike-trains demonstrated an average 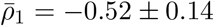 (mean ± s.d.) (*N* = 52). Nearly half of these fibers had *ρ*_1_ > −0.5 (*N* = 24) while the remaining fibers had *ρ*_1_ < −0.5 (*N* = 28). There was no clear relationship between the average firing rate of the neuron and its *ρ*_1_ (not shown). The second SCC (*ρ*_2_) between every other ISI (Fig. 2B) assumed positive (*N* = 36) or negative (*N* = 16) values with an average of *ρ*_2_ = 0.10 ± 0.18. The value assumed by *ρ*_2_ is linearly dependent on *ρ*_1_ with a positive *ρ*_2_ more likely if *ρ*_1_ < −0.5 (Fig. 2C). The linear relationship is described by the equation *ρ*_2_ = −1.18 *ρ*_1_ – 0.51, with a standard error (SE) of 0.008. This is close to the line *ρ*_2_ = – *ρ*_1_ – 0.5, the significance of which is discussed further below. Finally, the sum of the SCCs for each fiber taken over the first fifteen lags (excluding zeroth lag) is ∑_*k*_ *ρ_k_* = −0.475 ± 0.04, a number which, for most fibers, falls just short of −0.5 (Fig. 2D).

**Figure 2.**
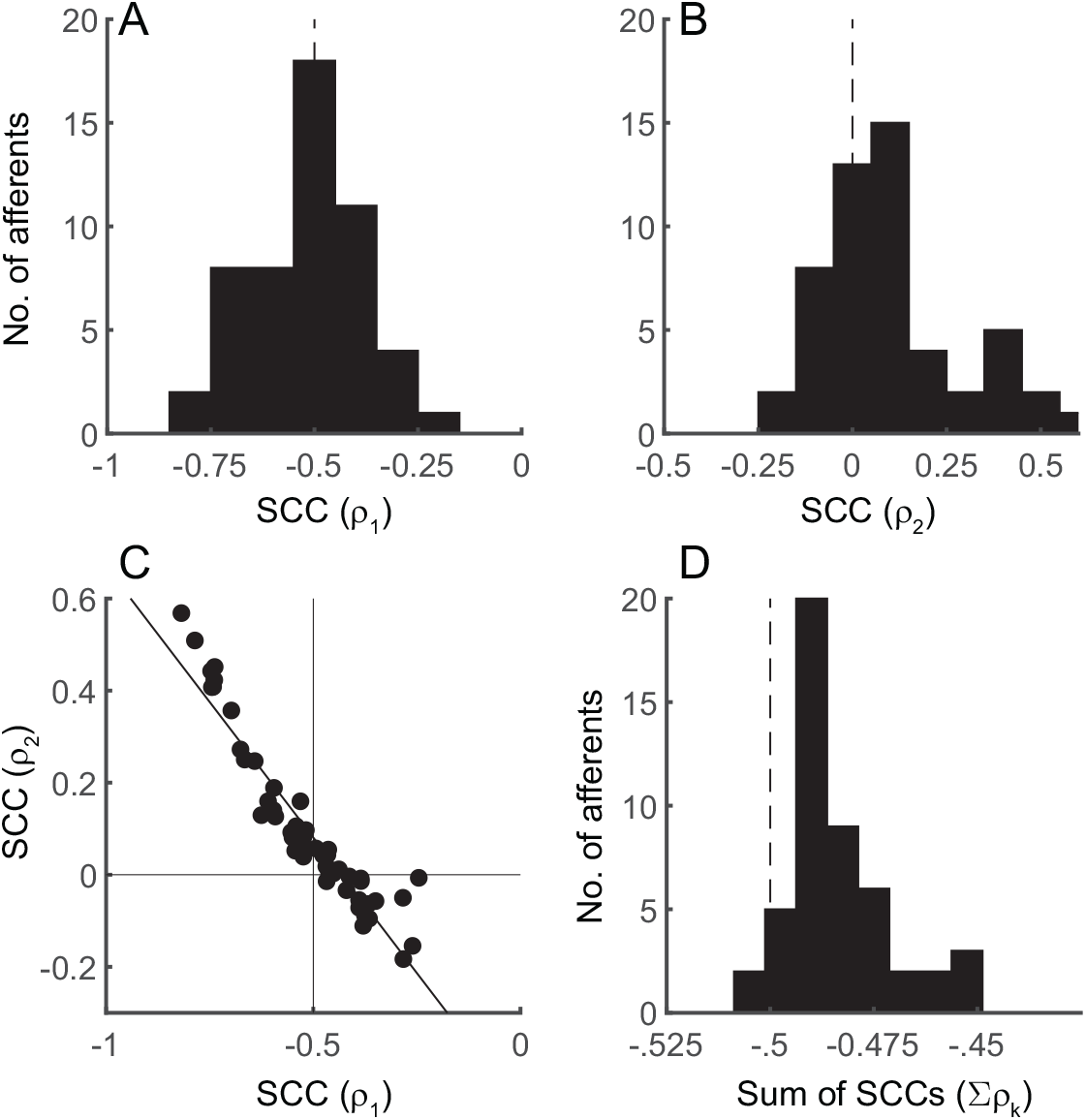
Population summaries of interspike interval (ISI) serial correlation coefficients (SCCs) for P-type primary afferent spike-trains (*N* = 52) [1]. A. Histogram of SCC between adjacent ISIs (*ρ*_1_), 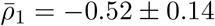 (mean ± s.d.), range −0.82 to −0.25 B. Histogram of SCC between every other ISI (*ρ*_2_), 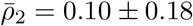, range −0.18 to 0.57. C. Anti-diagonal relationship between observed *ρ*_1_ (abscissa) and *ρ*_2_ (ordinate) (filled circles). Line describes best fit, *ρ*_2_ = −1.18*ρ*_1_ – 0.51 with SE = 0.008. D. Mean sum of SCCs for the population over 15 lags, ∑ *ρ_k_* = −0.475 ± 0.04. The population histograms in panels A and B were reported earlier with a different bin width (see [1], Figs. 7A and 7B therein, respectively).

### 3.2 Deterministic dynamic threshold model

In the simplest form of the dynamic threshold model [42], the spike-initiation threshold is a dynamic variable governed by two time-varying functions: the sub-threshold membrane potential *v*(*t*) and a time-varying or dynamic threshold *r*(*t*). In the sub-threshold regime *v*(*t*) < *r*(*t*). Most models (for e.g., [40]) assume that a spike is emitted when *v*(*t*) exceeds *r*(*t*) from below. To induce memory, the dynamic threshold is never reset (Fig. 1A) but suffers a jump in value whenever a spike is generated and then gradually relaxes to its quiescent value. This “jump and decay” is a fixed function which is called the dynamic threshold, and it usually takes the form *h*(*t*) = *A* exp (*-t/τ*) where *A* is the instantaneous jump in threshold following a spike at *t* = 0, and *τ* is the time-constant of relaxation. Between two spikes *t_k_* < *t* ≤ *t*_*k*+1_ the dynamic threshold is 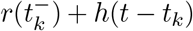 where 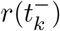 is the value assumed immediately before the spike at *t* = *t_k_*. It captures the sum over the history of spiking activity.

Most neuron models in the literature, including models incorporating a dynamic threshold, typically integrate the input using a low-pass filter (i.e., pass it through a leaky integrator) or a perfect integrator and then reset the membrane voltage to a constant *v_r_* following a spike (e.g., [17,37]). Some leaky integrators have been assumed to be non-resetting (e.g., [40]). In the form of the model considered in this work and earlier [42–44] the voltage *v*(*t*) is not integrated (i.e., there is no filtering), and it is not reset following a spike. These assumptions remove the effects of filtering in the soma and dendrites, and remove all the nonlinearities except for the spike-generator, which we assume generates a sequence of Dirac-delta functions.

For modeling spontaneous activity we consider a steady DC-level bias voltage *v* > 0. In its most general form, a spike is fired whenever *v* – *r*(*t*) = *γ*, where *γ* is a constant spike-threshold (Fig. 3A). Historically, and in much of the literature *γ* = 0 (for e.g., [17,37,40]). We use the more general form with a constant, non-zero *γ*. The specific value assumed by *γ* plays a role in optimal estimation of the stimulus [42,44] but does not influence serial correlation coefficients. This is addressed in the discussion. The major advantage of these simplifications is that they allow us to focus on the role of the dynamic threshold element *h* (*t*) in generating SCCs.

**Figure 3.**
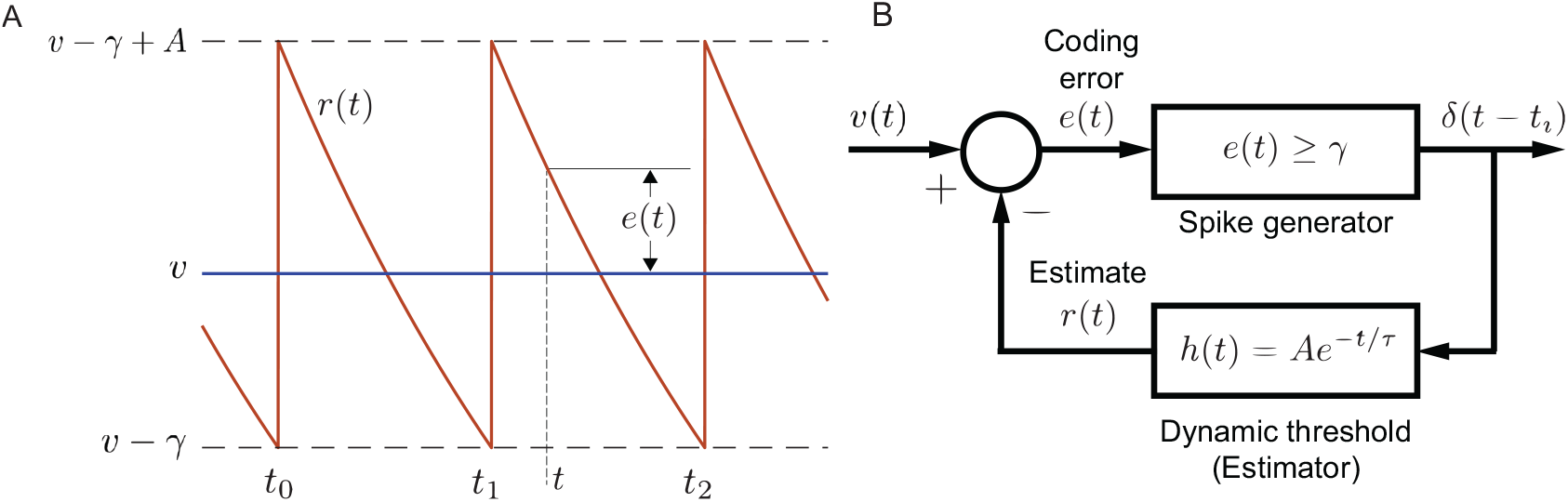
Schematic of neuron with deterministic dynamic threshold. A. *v* is a constant bias voltage that generates spontaneous activity, *r*(*t*) is a dynamic threshold, *γ* is a spike threshold such that a spike is emitted when *v* – *r*(*t*) = *γ*. Following a spike, the dynamic threshold suffers a jump of fixed magnitude *A.* The membrane voltage is non-resetting. Spike times are to, *t*_1_, *t*_2_,…, etc. Historically, the spike threshold was set to zero, i.e., *γ* = 0 and a spike is fired when *v* = *r*(*t*). Although it makes no difference to the results presented here, the more general form with *γ* is considered for reasons entered in the discussion. B. The spike encoder with dynamic threshold can be viewed as a feedback control loop where the spike-train *δ*(*t* – *t_i_*) is filtered by *h*(*t*) to generate the time-varying dynamic threshold *r*(*t*) which can be viewed as an estimate of the membrane voltage *v*(*t*). The comparator generates the estimation error *e*(*t*) = *v* – *r*(*t*) (see also Panel A). The estimation or coding error drives the spike generator, and a spike is fired whenever *e*(*t*) ≥ *γ*. The simplest form for the dynamic threshold (the estimation filter) *h*(*t*) mimics an RC membrane whose filter function is low-pass and given by *h*(*t*) = *A* exp (−*t/τ*). This is the form shown in panel A.

To make explicit the presence of memory, we note that the condition for firing the *k*^th^ spike at *t_k_* is met when

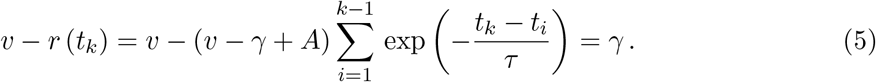

It is evident that memory builds up because of a summation over history of spiking activity. In the literature, a stochastic extension of this model is usually *v* – *r*(*t*) + *w* (*t*) = 0, where *w* is independent Gaussian noise. The spike encoder with dynamic threshold implicitly incorporates a feedback loop (Fig. 3B) and so a different view of the above model is to think of the dynamic threshold *r* (*t*) as an ongoing estimate of the membrane voltage *v* (*t*). Here, the dynamic threshold *h* (*t*) = *A* exp(−*t/τ*) acts as a linear low-pass filter. It filters the spike train to form an ongoing estimate *r* (*t*) of the voltage *v* (*t*). The instantaneous error in the estimate (the coding error) is then *e* (*t*) = *v* (*t*) – *r* (*t*) (Fig. 3B). When the error exceeds *γ* a spike is output and the threshold increases by *A* to reduce the error. The time-varying dynamic threshold tracks *v* (*t*) much like a home thermostat tracks a temperature set-point (Fig. 3A). Viewed in this way, it is the estimation error, and not the signal *v* (*t*), which directly drives the spike generator and determines when the next spike should be generated (Fig. 3B).

From Fig 3A we can approximate the exponential with a piece-wise linear equation when the ISI is small. If *t*_*i*–1_ and *t_i_*, *i* ≥ 1, are successive spike-times (Fig. 3B), then the time evolution of the dynamic threshold *r* (*t*) is given by

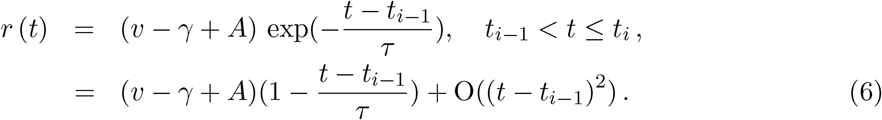

Note that *v, γ, A, τ* are constant, and so we can define *m* = (*v* – *γ* + *A*) /*τ* so that the slope of the decaying dynamic threshold is −*m*. The ISI can be obtained directly as (see Appendix for details)

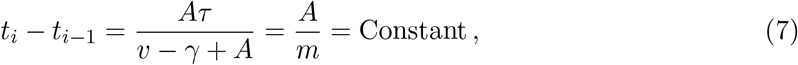

which is the deterministic firing rule for a spike generator with a constant, DC-level input voltage.

### 3.3 Stochastic extension of the dynamic threshold model

In the schematic shown in Fig. 3B, noise injected in the body of the feedback loop will reverberate around the loop and generate memory. Here we consider a noisy threshold *γ*. Subsequently we will provide some results for a noisy time-constant *τ* in the dynamic threshold element. Fig. 4 depicts the stochastic dynamic threshold model where the spike threshold (*γ*) is a stochastic process. All other aspects of the model are unchanged from the deterministic model (Fig. 3). Let *γ* be a discrete wide-sense stationary process with mean **E**[*γ*], discrete auto-correlation function **R**_*k*_ and power **E** [*γ*^2^] = **R**_0_. The spike threshold with additive noise assumes the random value *γ_i_*, *i* ≥ 1 immediately after the (*i* – 1)^th^ spike and remains constant in the time interval *t*_*i*–1_ < *t* ≤ *t_i_* (Fig 4A) [24,48]. Thus, the *i*^th^ spike is emitted when the error satisfies the condition

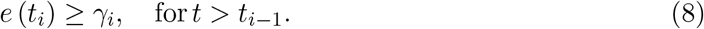

Subsequently the instantaneous value of the dynamic threshold jumps to a higher value specified by *v* – *γ_i_* + *A*, and the noisy spike threshold assumes a new value *γ*_*i*+1_. From Fig. 4A proceeding as before

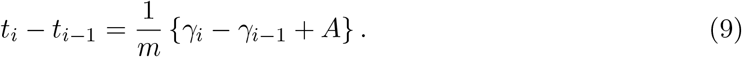

The mean ISI is therefore **E** [*t_i_* – *t*_*i*–1_] = *A/m* as in the deterministic case given by Eq. 7. From the assumption of wide-sense stationarity, the auto-correlation function **E** [*γ_i_ γ_j_*] can be written as **R** (*t_i_* – *t_j_*). This is a discrete auto-correlation function whose discrete nature is made clear in the following way (see also Appendix). Denote the mean of the *k*^th^-order interval by *T_k_* = **E**[*t*_*i+k*_ –*t_i_*], then the mean ISI is *T*_1_(= *A/m*), and further *T_k_* = *kT*_1_. A realization of the random variable *γ* is generated once every ISI and thus, **R** takes discrete values at successive multiples of the mean ISI, i.e., **R** (*T*_1_), **R** (*T*_2_), etc., and will be denoted by **R**_1_, **R**_2_, etc., respectively. As noted before, **R**_0_ is noise power. That is, we can write

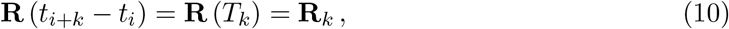

where *k* is the number of intervening ISIs. The general formula for serial correlation coefficients at lag *k* is well-known [49], and it is

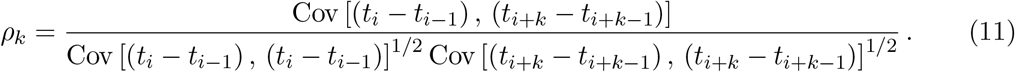

For a wide-sense stationary process the covariances are constant, so the subscript *i* can be dropped. From the relations and definitions in Eqs. 9, 10, and 11, and after some routine algebra (see Appendix), we obtain

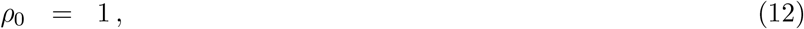

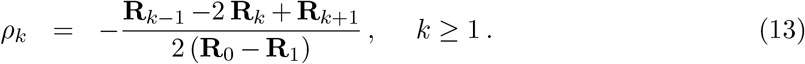

Further below we determine the **R**_*k*_ from experimental data. The serial-correlation coefficients given by Eqs. 12 and 13 are independent of the slope *m* of the decay rate of the dynamic threshold, and its gain *A*. Thus, for a constant input the observed correlation structure of the spike-train is determined solely by the noise added to the deterministic firing threshold *γ*.

**Figure 4.**
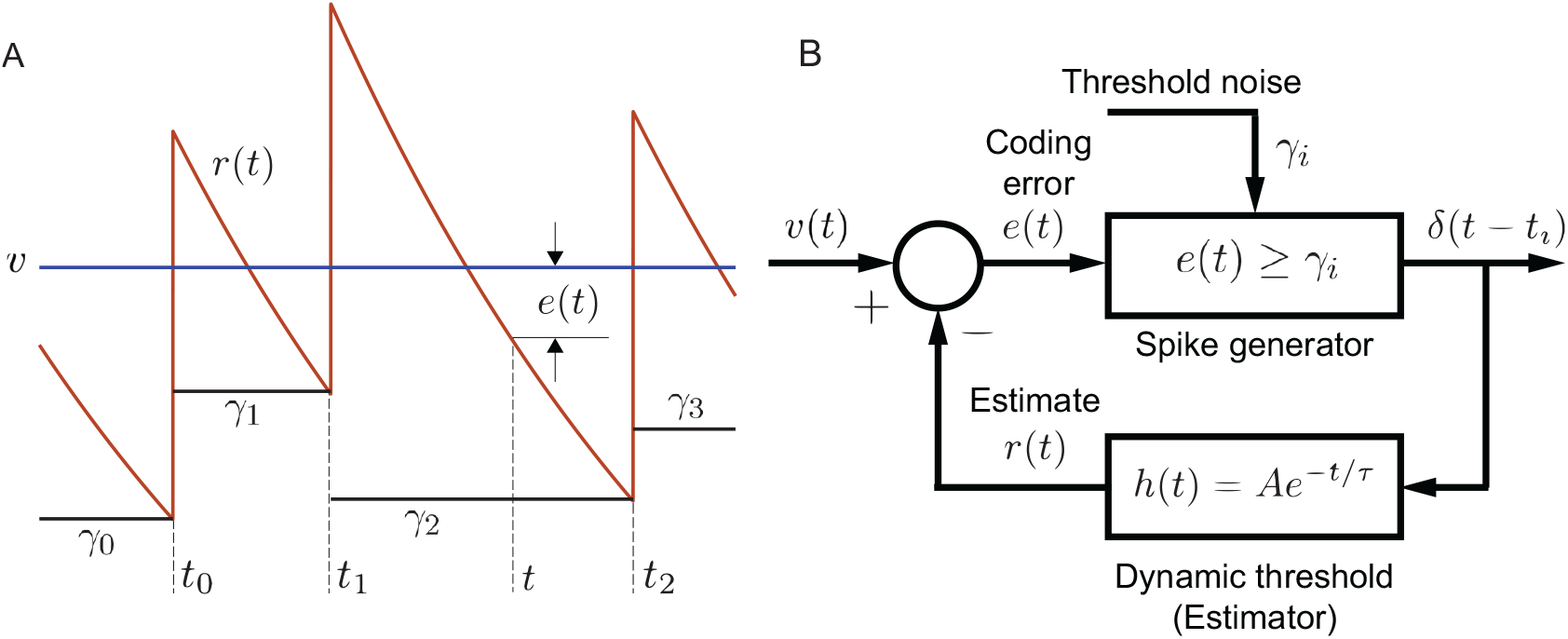
Schematic of a neuron with dynamic threshold and stochastic firing threshold. Panel descriptions follow Fig. 3 and only differences are noted. A. The spike threshold is *γ_i_* = *v* – *r* (*t*) where *γ_i_* is a random value generated at 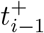, and held constant until the next spike at *t_i_*. The discrete noise sequence *γ_i_* is generated by a discrete wide-sense stationary process with mean *γ* and unknown discrete auto-correlation function **R**_*k*_. The goal is to determine **R**_*k*_ which will generate a prescribed sequence of ISI serial correlation coefficients *ρ_k_*. To reduce clutter, the spike threshold *v* – *γ_i_* is depicted as *γ_i_*. B. Block diagram showing the stochastic modification of the spike threshold. Symbols and additional description as in text and Fig. 3.

#### 3.3.1 Limiting sum of SCCs and power spectra

We make the assumption that the process *γ* is aperiodic and the auto-correlation function **R**_*N*_ → 0 when *N* → ∞. That is, the noise process decorrelates over long time-scales. From this assumption and Eq. 13 it follows that (see Appendix)

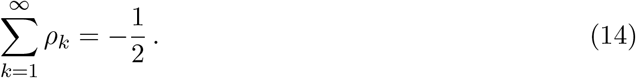

This is the limiting sum of ISI serial correlation coefficients. Let the mean and variance of ISIs be denoted by *T*_1_ and *V*_1_, respectively, and the coefficient of variation of ISIs as 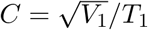. If *P* (*ω*) is the power spectrum of the spike train, then the DC-component of the power is given by [49]

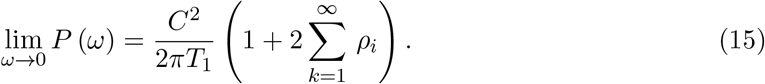

Introducing Eq. 14 into Eq. 15, we obtain

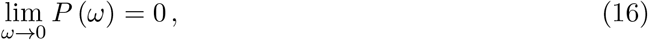

yielding a perfect DC block.

The limiting sum of SCCs from experimental spike trains, with the sum calculated up to 15 lags are depicted in Fig. 2D. The population of afferents demonstrated a range of limiting sums with mean sum of −0.475 ± 0.04. The limiting sum was less than −0.5 in only two afferents, where the sums were estimated as —0.505 and −0.502.

#### 3.3.2 Predicting Type I and Type II serial correlation coefficients

In the relationships for the SCCs given by Eqs. 12 and 13, the noise process that drives the spike generator has an unknown correlation function **R** which must be determined so as to predict experimentally observed SCCs. Here, we are interested in identifying the simplest process that can satisfactorily capture observed SCCs. Consider a Gauss-Markov, i.e., Ornstein-Uhlenbeck process which relaxes as exp(−*t/τ_γ_*) with relaxation time *τ_γ_*, where *γ* signifies the spike threshold. We are interested in a discrete-time realization of the process. Thus, we define a parameter *a* = exp (−*T*_1_/*τ_γ_*) < 1, where *T*_1_ is the mean ISI. We can now define two first-order autoregressive (AR) processes that will generate Type I and Type II SCCs, respectively.

##### Type I serial correlation coefficients

Type I afferent spiking activity demonstrates serial correlations where −0.5 < *ρ*_1_ < 0, and subsequent *ρ*_k_ are negative and diminish to 0 with increasing *k* (Fig. 1A). These spike trains have a unimodal ISI and do not display bursting activity. They can be generated from the ansatz, a first-order AR process

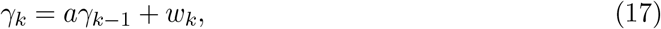

where *γ_k_* is the noise added to the threshold at the *k*^th^ spike (Fig. 4), *a* is as defined above, and *w_k_* is white noise input to the AR process. The output *γ* is wide sense stationary and its discrete autocorrelation function **R**_*k*_ is

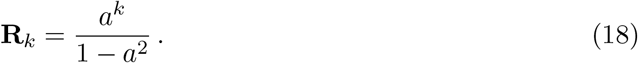

Eqs. 13 and 18 yield the geometric series

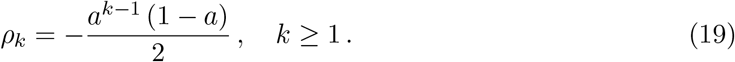

From Eq. 19, and noting that 0 < *a* < 1, we conclude that *ρ*_1_ > −0.5, *ρ*_k_ < 0 for all *k*, |*ρ*_*k*–1_| > |*ρ_k_*|, and *ρ*_k_ → 0 as *k* → ∞. This is the observed Type I pattern of SCCs. Further, summing the geometric series Eq. 19 yields ∑_*k*≥1_ *ρ*_k_ = −0.5 as stated in Eq. 14. The AR parameter *a* can be estimated from experimentally determined SCCs. From Eq. 19 this is

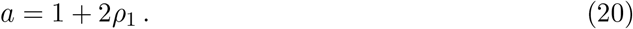

In practice, a better estimate is often obtained from the ratio *a* = *ρ*_2_/*ρ*_1_, where *ρ_k_* are available from experimental data.

We model a Type I P-type primary electrosensory afferent depicted in Fig. 1A using a noisy threshold with correlation function **R**_*k*_ specified by Eq. 18 and shown in Fig. 5A. The dynamic threshold parameters were determined from the experimental data, and tuned so that they matched the afferent SCCs, ISI, and joint distributions (Fig. 6). By design, the noise process is first-order autoregressive, and this is reflected in the partial autocorrelation function calculated from the noise samples (Fig. 5D). The top row (Figs. 6A-C) depicts the ISI distribution, joint ISI distribution, and the serial correlation coefficients, respectively. Type I spike trains do not display bursting activity and their ISI distribution is unimodal. The bottom row (Figs. 6D-F) shows data from a matched model using noise correlation function given by Eq. 18. SCCs of adjacent ISIs *ρ*_1_ are −0.39 (data) and −0.40 (model). The mean sum of SCCs are −0.48 (data) and −0.5 (model). Thus, the observed pattern of Type I SCCs is reproduced.

**Figure 5.**
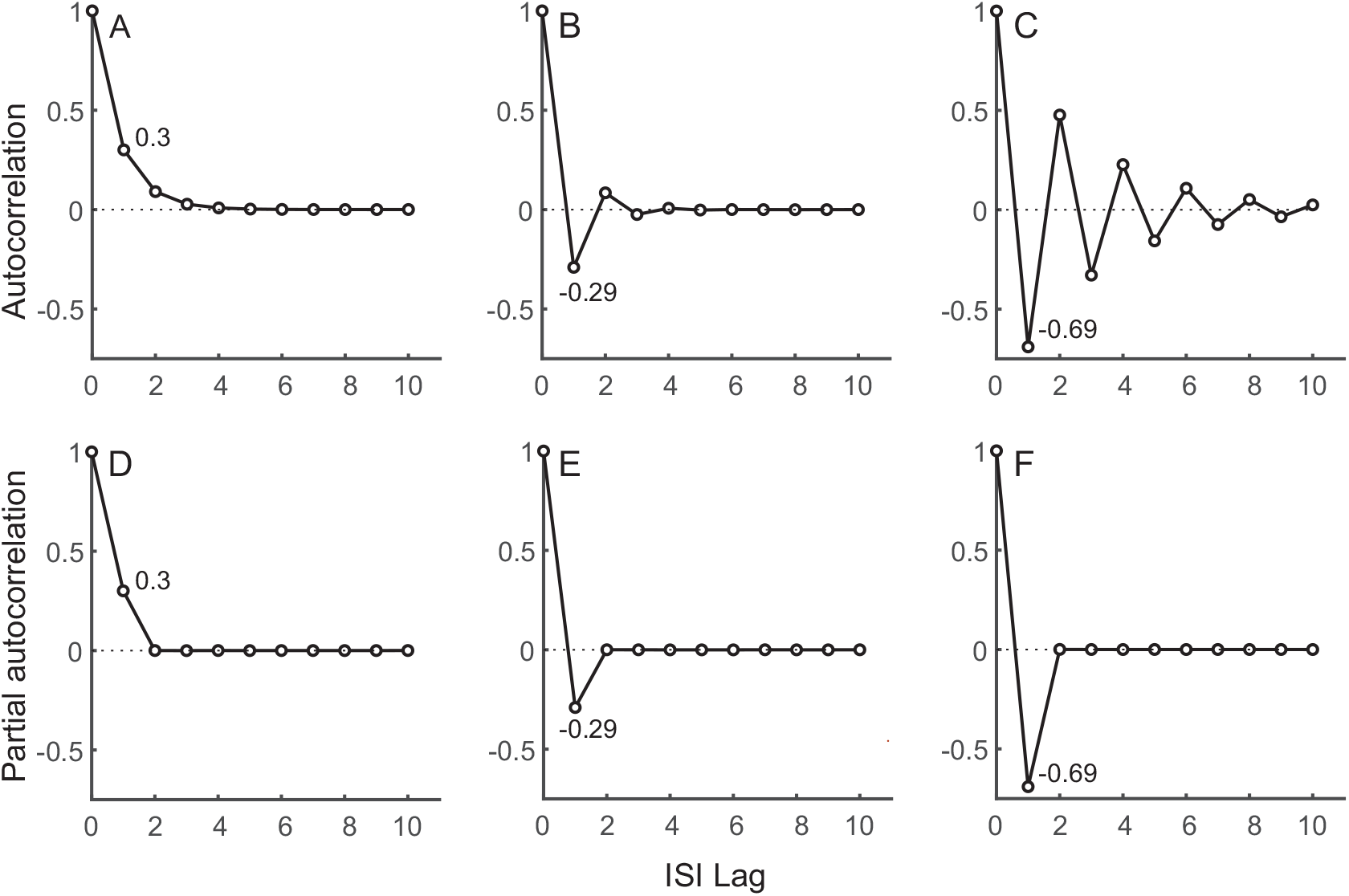
Threshold noise: Normalized autocorrelation (top row) and partial autocorrelation functions (bottom row) of threshold noise. A. Type I SCC matched to the afferent depicted in Fig. 1A with no bursting activity. B. Type II SCC from an afferent with moderate bursting activity. C. Type II SCCs matched to the afferent depicted in Fig. 1B with strong bursting activity. Autocorrelation function that matches Type III SCCs will be zero for all non-zero lags (not shown). Panels D-F show partial autocorrelation functions of the corresponding noise process shown in panels A-C, respectively. The partial autocorrelations are non-zero for the first lag, and zero for all lags *k* > 1. This is because the noise generator is a first-order autoregressive (AR) process.

**Figure 6.**
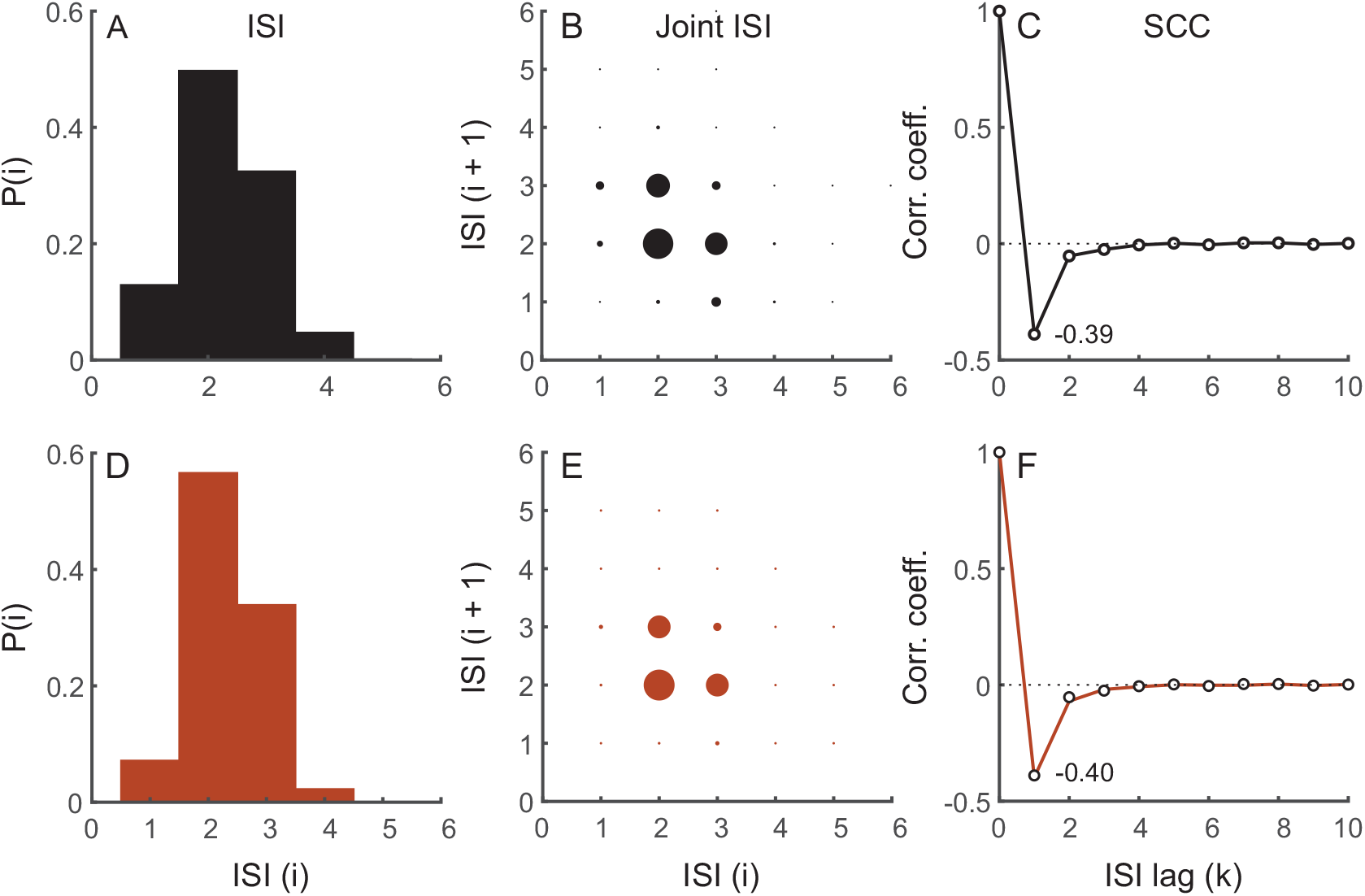
Type I serial correlation coefficients (SCCs). Top row depicts experimental spike-train from a P-type primary electrosensory afferent. A. The spike train has a unimodal interspike interval distribution (ISI) and does not display bursting activity. Abscissa is ISI in multiples of electric organ discharge (EOD) period and ordinate is probability. B. Joint interspike interval distribution showing probability of observing successive intervals ISI (*i*) (abscissa) and ISI(*i* + 1) (ordinate). The size of the circle is proportional to the probability of jointly observing the adjacent ISIs, i.e., *P* (*i, i* + 1). C. Serial correlation coefficients (ordinate) as a function of lag measured in multiples of mean ISI (abscissa). Spike-train sampled at EOD period. SCCs for this afferent are shown in Fig. 1A at a different sampling rate (see Methods). Bottom row depicts results for matched model using colored noise with correlation function given by Eq. 18 and shown in Fig. 5A. Panel descriptions as in top row, except in F where open circles denote experimental data taken from C. EOD period: 1.06 ms, mean ISI: 2.42 ms. To generate model results, *v* = 1.845 V, adaptive threshold parameters: *A* = 0.15 and *τ* = 30 ms, AR parameter: *a* = 0.4 (*τ_γ_* = 2.09 ms), and **R**_0_ = 1.07 × 10^-3^ V^2^ (SNR = 35 dB).

##### Type II serial correlation coefficients

Type II afferent spiking activity demonstrates serial correlations where *ρ*_1_ < −0.5 and successive *ρ_k_* alternate between positive (*ρ*_2*k*_) and negative (*ρ*_2*k*+1_) values, are progressively damped, and diminish to zero (Fig. 1B). These spike trains have bimodal ISIs and display bursting activity. Proceeding as before, they can be generated from the ansatz, a first-order AR process

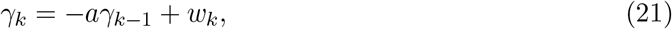

with discrete autocorrelation function

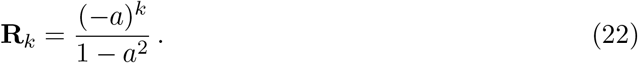

Eqs. 13 and 22 yield the geometric series

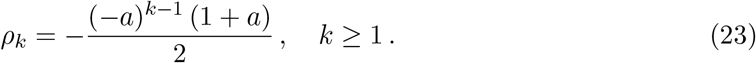

From Eq. 23, and noting that 0 < *a* < 1, we conclude that *ρ*_1_ < −0.5, *ρ*_2*k*_ > 0, *ρ*_2*k*+1_ < 0, |*ρ*_2*k*_| > |*ρ*_2*k*+1_|, and *ρ_k_* → 0 as *k* → ∞. This is the observed Type II pattern of SCCs. Further, summing the geometric series (Eq. 23) yields ∑_*k*≥1_ *ρ_k_* = −0.5 as stated in Eq. 14. The AR parameter a can be estimated from experimentally determined SCCs. From Eq. 23 this is

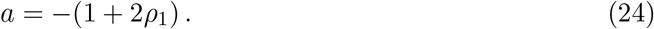

As noted for Type I SCCs a better estimate is often obtained from the ratio *a* = −*ρ*_2_/*ρ*_1_. Finally, note that Type II serial correlations can be obtained from Type I serial correlations by replacing a in Eqs. 18, 19, and 20 with −*a*, to obtain Eqs. 22, 23, and 24, respectively.

We model Type II P-type primary electrosensory afferents using a noisy threshold with correlation function **R**_*k*_ specified by Eq. 22. The noise correlation function and dynamic threshold parameters were determined from the experimental data, and tuned so that they matched the afferent SCCs, ISI, and joint distributions. We demonstrate with two examples: i) moderate bursting activity and ii) strong bursting activity. The moderately bursting neuron has a broad bimodal ISI distribution (Fig. 7A-C), and its ISI and joint ISI distributions, and SCCs are captured by the matched model (Fig. 7D-F). The noise correlation function for generating the model spike-train is depicted in Fig. 5B, and the noise partial correlation function in Fig. 5E. SCCs of adjacent ISI *ρ*_1_ are −0.59 (data) and −0.62 (model). The mean sum of SCCs are −0.49 (data) and −0.5 (model). The strongly bursting neuron has a well-defined bimodal distribution (Fig. 8A-C), and its ISI and joint ISI distributions, and SCCs are captured by the the matched model (Fig. 8D-F). The noise correlation function for generating the model spike-train is depicted in (Fig. 5C), and the noise partial correlation function in Fig. 5F. SCCs of adjacent ISI *ρ*_1_ are −0.7 (data) and −0.7 (model). The mean sum of SCCs are −0.49 (data) and −0.5 (model). Thus, in both cases (Figs. 7 and 8), the observed patterns of Type II SCCs are reproduced. All observed afferent spike trains were either Type I or Type II.

**Figure 7.**
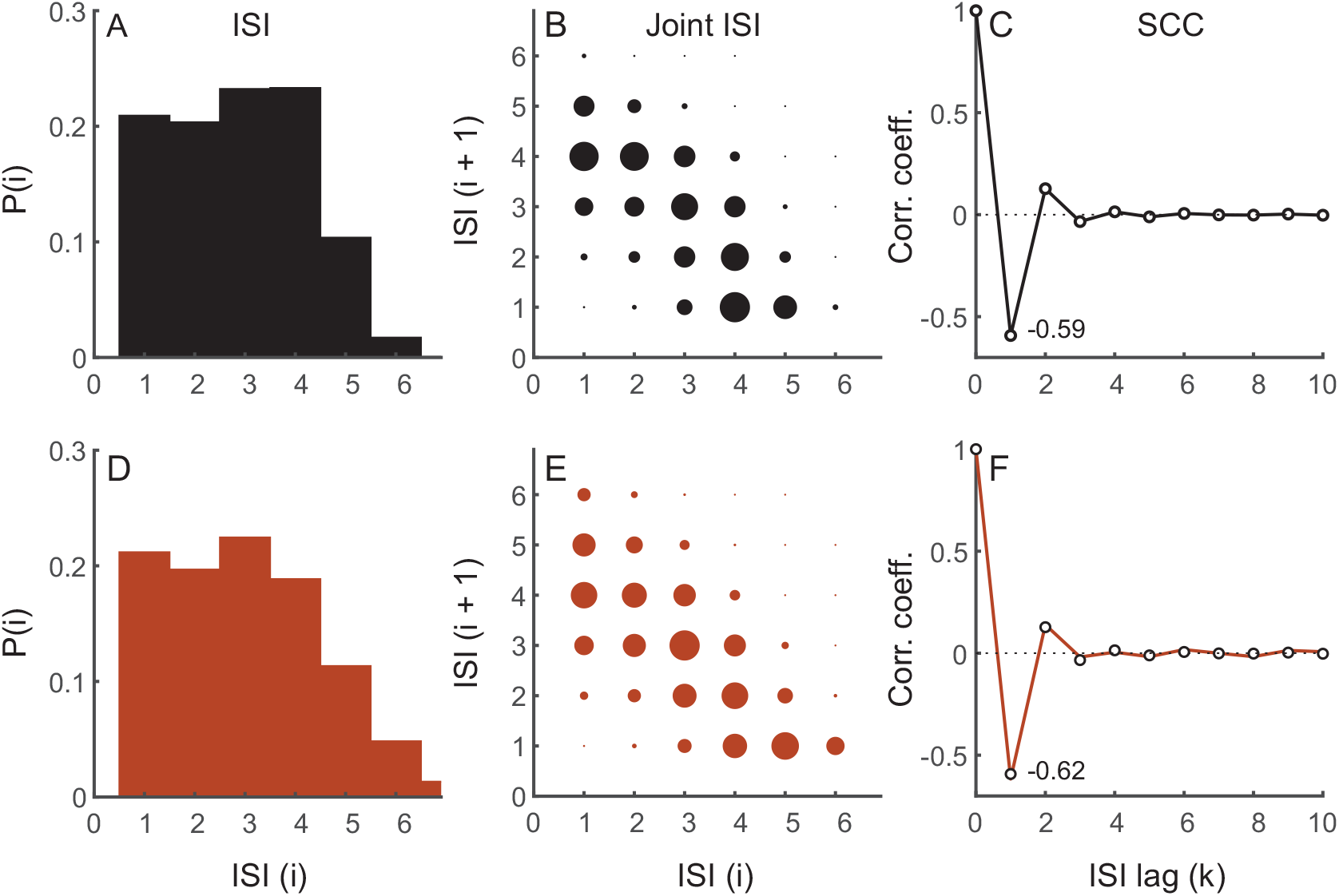
Type II serial correlation coefficients (SCCs) with moderate bursting activity. Description is identical to Fig. 6. Top row depicts experimental spike train from a P-type primary electrosensory afferent. Bottom row depicts results for matched model using using colored noise with correlation function given by Eq. 22 and shown in Fig. 5B. EOD period: 1.31 ms, mean ISI: 3.77 ms. To generate model results, *v* = 1.845 V, adaptive threshold parameters: *A* = 0.28 and *τ* = 26 ms, AR parameter: *a* = 0.29 (*τ_γ_* = 3.18 ms), and **R**_0_ = 9.22 × 10^-3^ *V*^2^ (SNR = 26 dB).

**Figure 8.**
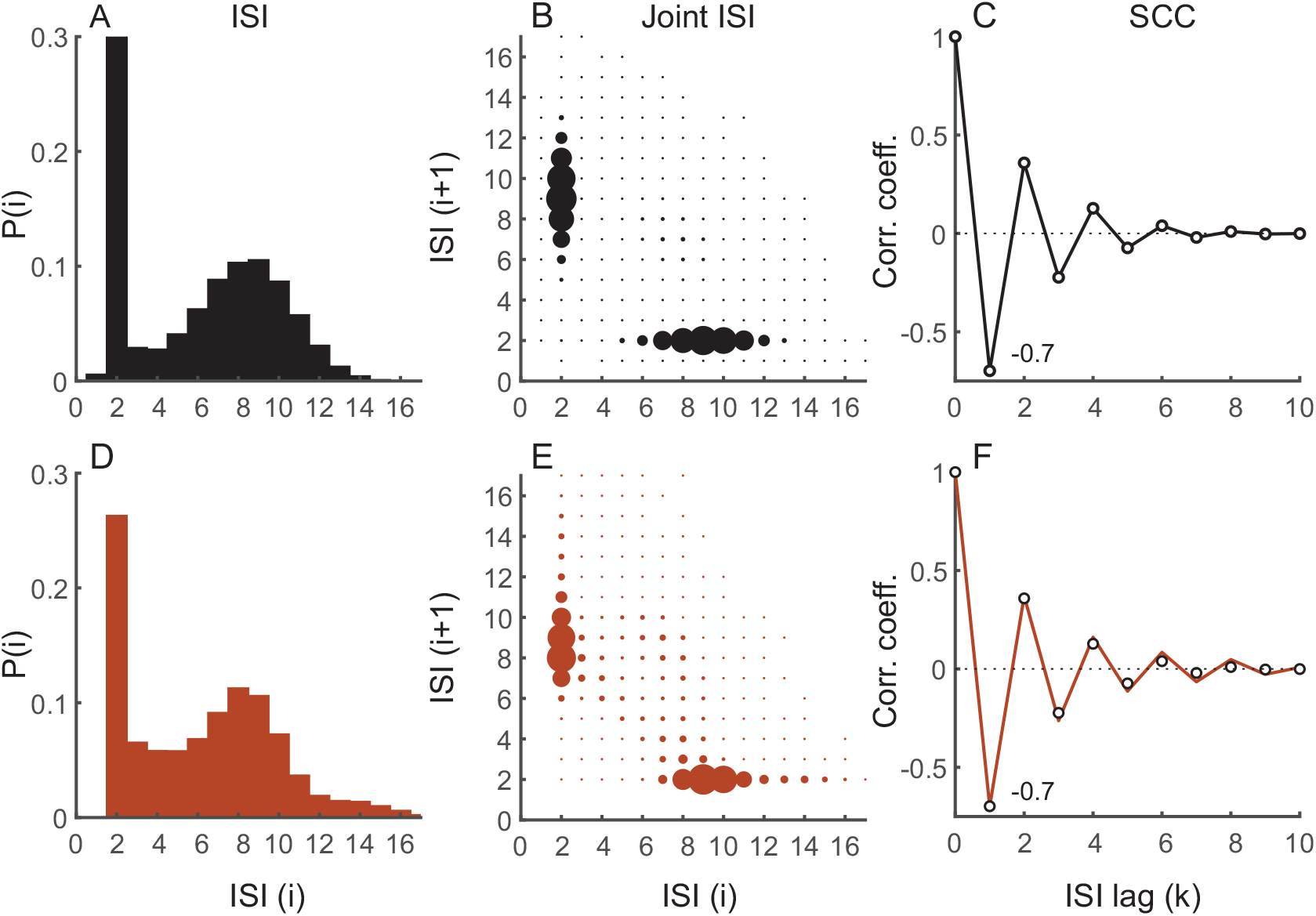
Type II serial correlation coefficients (SCCs) with strong bursting activity. Description is identical to Fig. 6. Top row depicts experimental spike train from a P-type primary electrosensory afferent. SCCs for this afferent are shown in Fig. 1B at a different sampling rate (see Methods). Bottom row depicts results for matched model using using colored noise with correlation function given by Eq. 22 and shown in Fig. 5C. EOD period: 1.04 ms, mean ISI: 6.04 ms. To generate model results, *v* = 1.845 V, adaptive threshold parameters: *A* = 0.19 and *τ* = 60 ms, AR parameter: *a* = 0.69 (*τ_γ_* = 16.89 ms), and **R**_0_ = 6.4 × 10^-3^ V^2^ (SNR = 27 dB).

We mention in passing that the noise correlation functions depicted in Fig. 5A-C should not be confused with the SCCs depicted in panels C and F in Figs. 6, 7, and 8.

##### Type III serial correlation coefficients

Type III afferent spiking activity demonstrates serial correlations where *ρ*_1_ = −0.5 and all *ρ_k_* = 0 for *k* ≥ 2. This is a degenerate case, resulting in a singleton with a unique set of SCCs. Such spike trains can be generated by a process where spike threshold noise is uncorrelated, and hence white. In this case, 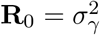 is noise power, and **R**_*k*_ = 0 for *k* ≥ 1. We see immediately from Eqs. 12 and 13 that the *ρ_k_* have the prescribed form. Further, we note that trivially ∑_*k*≥1_ *ρ_k_* = −0.5 as stated in Eq. 14. We mention in passing, and for later discussion, that white noise added to the spike threshold is equivalent to setting *a* = 0 (*τ_γ_* = 0) in the first-order AR process defined in Eq. 17. A spike-train with exactly Type III SCCs has not been observed in the experimental data presented here.

#### 3.3.3 Partial autocorrelation functions

Partial correlations were estimated from experimental data and the modeled spike trains using Eqs. 2, 3, and 4. These are shown in Figs. 9A, 9B, and 9C for the afferents shown in Figs. 6, 7, and 8, respectively. By definition the first partial autocorrelation *ϕ*_1,1_ = *ρ*_1_. In contrast to the SCCs where correlation coefficients *ρ_k_* could be positive or negative for *k* ≥ 2, all partial autocorrelations are negative irrespective of the type of afferent.

**Figure 9.**
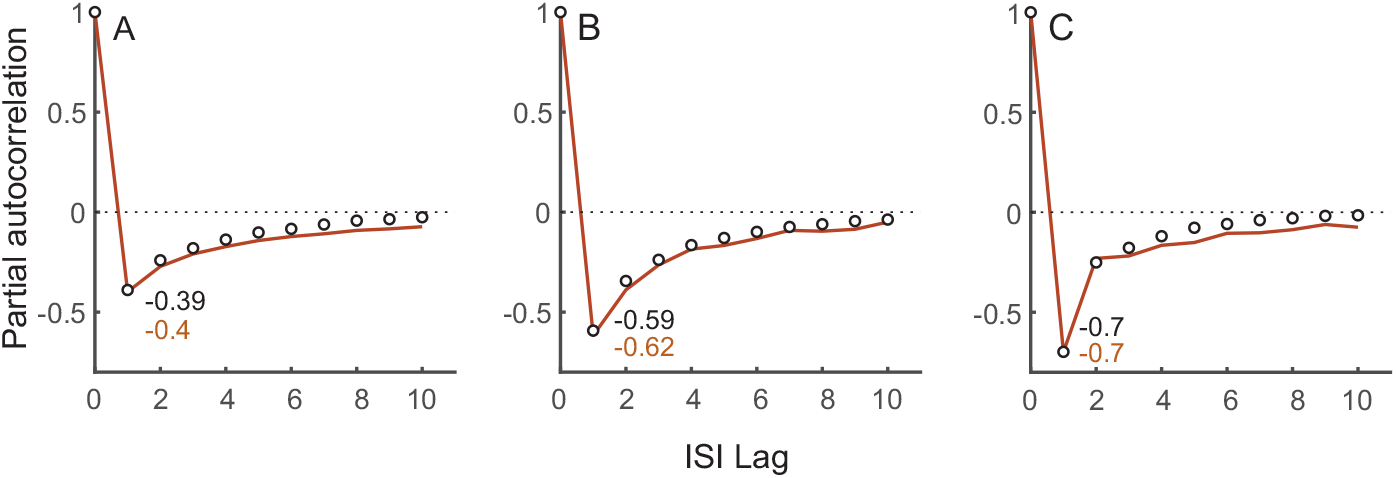
Partial autocorrelation functions estimated from afferent ISI sequences (open circles) and models (red line) for three afferents shown earlier: A. Type I, non-bursting (Fig. 6F). B. Type II, moderately bursting (Fig. 7F). C. Type II, strongly bursting (Fig. 8F). The first partial correlation coefficient *ϕ*_1,1_ is reported in each panel (data: black text, model: red text), and is the same as *ρ*_1_ (Eq. 4). Afferent and model spike-trains likely match an autoregressive moving average (ARMA) model of unknown order.

#### 3.3.4 Dynamic threshold with a random time-constant

The dynamic threshold has three parameters *A, γ*, and *τ*. We have so far described a stochastic model based on a noisy *γ* which is formally equivalent to a noisy *A*. We now consider a stochastic dynamic threshold model based on a noisy time-constant *τ*. We can transform the adaptive threshold model with a random spike threshold from the previous section and Fig. 4 so that the time-constant of the adaptive threshold filter *h* (*t*) is a random variable with mean *τ* (Fig. 10). From Eq. 9 the random variate *γ_i_* — *γ_i_*-1 which appears in the time-base can be transformed into the random variate mi which is the slope of the linearized adaptive threshold (green line, Fig. 10). From

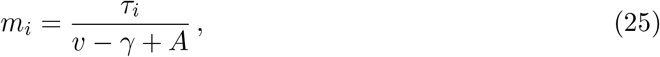

we obtain

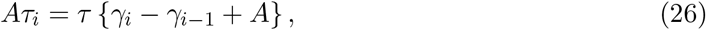

and so, **E** [*τ_i_*] = *τ*. As noted, this is the mean value of the dynamic threshold filter time-constant. It is immediately apparent from Eqs. 9 and 26 that the covariance of the sequence τi (sampled at the ISIs) is the same as the covariance of the ISI sequence up to a scale-factor, and therefore the serial correlations of *τ* (*ρ_τ,k_*, *k* ≥ 0) are the same as the serial correlations of the ISIs. Therefore (see Appendix for details)

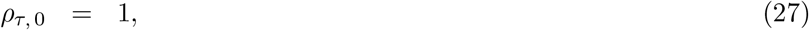

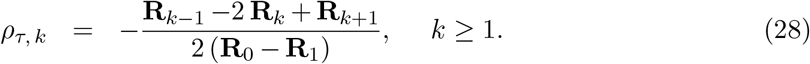

The right side of the above equations, Eqs. 27 and 28, are the same as Eqs. 12 and 13, the expressions for the SCCs of a spike train generated with a noisy adaptive threshold. Thus, the random filter time-constants have the same serial correlation coefficients as the interspike intervals, *ρ_τ,k_* = *ρ_k_*. This is a “pass through” effect where the correlations observed in the time-constant are directly reflected in the ISS correlations.

**Figure 10.**
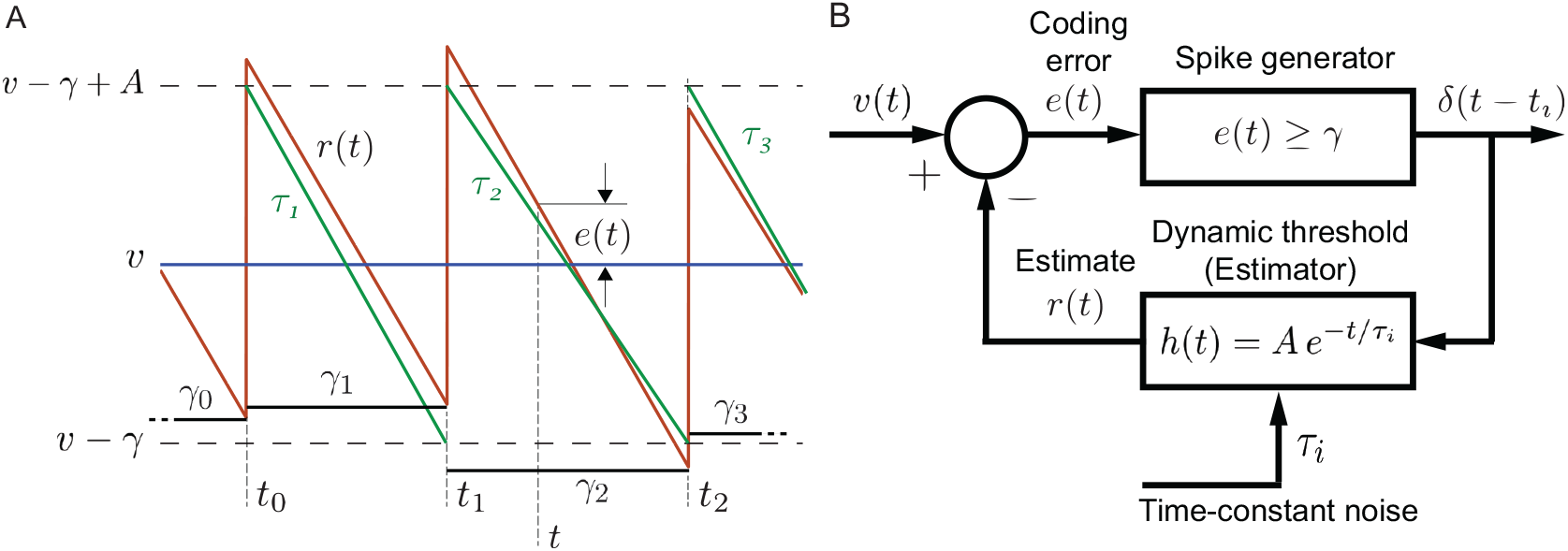
Dynamic threshold with a stochastic filter time-constant. B. Geometry of the spiking mechanism. The input is a deterministic piece-wise constant signal (*v*, here shown to be DC-valued). Two equivalent spike-generation processes are possible: Case (i) The spike threshold *γ* is noisy while the dynamic threshold filter *h*(*t*) has fixed time-constant *τ*. This is the same as Fig. 4A. The spike threshold is a constant term *v* – *γ* plus a random perturbation *x_i_* generated at 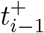, and held constant until the next spike at *t_i_* (black). The time-varying dynamic threshold *h*(*t*) is shown in red. Case (ii) The function *h*(*t*) shown in green has a random time-constant with mean *τ* while the spike threshold is *v* – *γ*, and is constant. Threshold noise *x_i_* from Case (i) can be transformed into a noisy dynamic threshold time-constant *τ_i_* which is fixed between spikes, and takes correlated values over successive ISIs. Spike-times for both cases are identical. B. Block diagram illustrating the feedback from the estimator that generates optimal spike times. The time-constant *τ* is a random variable that is constant between spikes, and is correlated over interspike intervals (ISIs). It is of interest to ask what is the relationship between the correlation functions in the two cases. Symbols and additional description as in text.

In summary, a noisy threshold *γ* or a noisy filter time-constant *τ* can be used to generate spike trains which have prescribed ISI SCCs. We have generated spike trains using a noisy threshold and will not duplicate the results for a noisy time-constant.

## 4 Discussion

### 4.1 Experimental observations

All experimentally observed P-type spike-trains from *Apteronotus leptorhynchus* demonstrated negative correlations between adjacent ISIs (Figs. 1, 2A). Thus, negative SCCs between adjacent ISIs may be an obligatory feature of neural coding, at least in this species. The implications for coding are discussed further below. A broad experimental observation is the roughly equal division of spike-trains into units with *ρ*_1_ > −0.5 (non-bursting or Type I units, Fig. 1A, *N* = 24) and units with *ρ*_1_ < −0.5 (bursting or Type II units, Fig. 1C, *N* = 28). Using a different method of classification, Xu *et al.* [50] reported 31% of 117 units as bursting. There is no clear relationship between the mean firing rate of a unit and its *ρ*_1_ (not shown). Further, for Type I units *ρ*_2_ < 0, whereas for Type II units *ρ*_2_ > 0 (Fig. 2C). The former give rise to over-damped SCCs which remain negative and diminish to zero, while the latter give rise to under-damped or oscillatory SCCs which also diminish to zero. The dependence of *ρ*_2_ on *ρ*_1_ is linear for the most part and follows the equation *ρ*_2_ = −1.18*ρ*_1_ – 0.51 (Fig. 2C). The limiting sum of SCCs ∑_*k*≥1_ *ρ_k_* is close to −0.5 irrespective of the type of SCC pattern (Fig. 2D) [19]. With the exception of two afferent fibers, the sum of SCCs for the fibers were never smaller than −0.5, i.e., it was almost always true that ∑_*k*≥1_ *ρ_k_* ≥ −0.5. In two cases the sums were less than the limit (−0.505 and −0.502), possibly due to estimation error. Although the dominant SCCs are *ρ*_1_ and *ρ*_2_, their sum *ρ*_1_ + *ρ*_2_ is not close to −0.5 (i.e., *ρ*_2_ ≠ – *ρ*_1_ – 0.5) and deviates, as stated above, with a linear relationship which follows *ρ*_2_ = −1.18*ρ*_1_ – 0.51. Thus, more terms (lags) are needed to bring the sum of SCCs close to the limiting value, and this results in significant correlations extending over multiple lags (time-scales). As discussed in an earlier work [1], long-short (short-long) ISIs create memory in the spike-train and keep track of the deviations of successive ISIs from the mean ISI (*T*_1_). These deviations are a series of “credits” and “debits” which may not balance over adjacent ISIs, but will eventually balance so that a given observation of *k* successive ISIs returns to the mean with *t_k_* – *t*_0_ ≈ *kT*_1_. Such a process will exhibit long-range dependencies that may not be captured by SCCs.

That the dependencies may extend over multiple ISIs is confirmed from an analysis of joint dependencies of intervals extending to high orders [1,6]. All 52 units in *Apteronotus leptorhynchus* were at least second-order Markov processes, with about half (*N* = 24) being fifth-order or higher [1]. Further, SCCs were not correctly predicted when only the adjacent ISI dependencies were preserved, i.e., were considered to be first-order Markov [1,51,52]. Indeed, an examination of the sequence of SCCs provides no indication of the extent of memory. For instance, short-duration correlations do not necessarily imply that ISI dependencies are limited to fewer adjacent intervals. Long-duration dependencies may be present even when the correlation time is short [6]. Conversely, a first-order Markov process produces a ringing in the serial correlogram 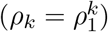 that can continue for ISIs much longer than two adjacent ISIs [49,53]. In fact, for some P-type electrosensory afferent spike trains, the observed ISIs exhibited SCCs whose magnitudes were smaller than the SCCs for the matched first-order Markov model even though the experimental data was at least second-order or higher (see Fig. 8, [1]).

The stochastic process which generates ISIs may be more complex than a simple autoregressive (AR) or moving average (MA) process because afferent ISI correlation functions (Figs. 6F, 7F, and 8F) and partial autocorrelation functions (Fig. 9) are infinite in duration (see Methods for a distinction between the two functions) [47]. This suggests that a more general ARMA process may be responsible for the generation of ISIs. The source of the mix of AR and MA processes is discussed further below when we consider the generating model.

### 4.2 The dynamic threshold model

The experimental observations and dependency analyses motivated us to ask whether we could develop a stochastic model to reproduce a prescribed sequence of SCCs *ρ*_1_, *ρ*_2_,…, *ρ_k_*. We adopt a widely used and physiologically plausible model with a time-varying, i.e., dynamic, threshold. This is a simple model without much complexity, and has few parameters. The model allows us to probe patterns of SCCs as a function of these parameters. Further, it tests the extent to which we can describe experimental data with a simple model. Dynamic threshold models typically have three components: 1) dynamics of membrane voltage *v*(*t*) in response to an input signal, 2) a spike or impulse generator, and 3) a time-varying dynamic threshold *r*(*t*) which is elevated following a spike and subsequently relaxes monotonically. A spike is fired when the membrane voltage meets the (relaxing) threshold, thus forming a feedback loop (Fig. 3B). Models of adaptation with an adaptation current are broadly similar but instead of altering the threshold directly, they use an outward current to induce refractoriness (see for e.g., [26,27,37]). In many models, the feedback loop is implicitly defined (as in conductance-based models), but it may also be explicitly defined as is done here so that *v*(*t*) – *r*(*t*) reaches a fixed spike-initiation threshold. This threshold is usually taken to be zero, i.e., *v*(*t*) = *r*(*t*) [40], but here we assume it to be a non-zero value (*γ* = *A*/2) following [42]. This choice does not alter model behavior or the analyses presented here (see Methods) but has relevance to optimal coding and is discussed further below.,

Models that do not utilize a dynamic threshold or an adaptation current are not discussed here because they are outside the scope of this work. The reader may consult [3,46,54,55] for more details and references to such models. Most dynamic threshold models that address negative correlations assume some form of perfect or leaky integrate-and-fire dynamics for the first two components listed above [17,23,27,34,35], and an exponentially decaying dynamic threshold without spike reset. Noise is added to the time-varying threshold or to the input current and this results in negative serial correlations. Model complexity has been a major drawback in determining the precise role of noise in shaping ISI correlations (see for instance observations made in [23, 56]). Resets, hard refractory periods, sub-threshold dynamics due to synaptic filtering, and sometimes multiple sources of noise obscure the effects of signal propagation through the system and obscure signal dependencies. Thus with few exceptions (see below), dynamic threshold models and models with adaptation currents, have been qualitative. They demonstrate some features of experimentally observed ISI distributions, and at best correlations between adjacent ISIs (i.e., *ρ*_1_) [32,35,45]. On the other hand, reduced model complexity can result in a lack of biophysical plausibility. Thus a judicious choice of models should expose desired mechanisms while retaining enough important features of the phenomena.

### 4.3 Model results

In recent years deterministic dynamic threshold models with an exponential kernel have been used to predict spike-times from cortical and peripheral neurons [40, 42,57] (see [41] for an early review) and predict peri-stimulus time histograms [42, 44] with good accuracy. Capturing spike-times accurately is perhaps the first requirement in our analysis, and this gives confidence that the model may tell us something about ISI correlations. We eliminated sub-threshold dynamics and resets so that there is only one nonlinear element, the spike generator [42, 44]. These are not serious restrictions, and they make the analysis tractable. We follow the usual practice of representing the dynamic threshold element with an exponential decay with time-constant *τ*. The absence of reset implies that the time-varying dynamic threshold which carries memory is a simple convolution of the spike train with an exponential kernel *Ae*^−*t/τ*^ (Fig. 3B). We inject noise precisely in one of two places, either to perturb the spike threshold *γ* (Fig. 4) or perturb the time-constant *τ* (Fig. 10). The two forms of perturbation are formally equivalent (see Appendix). We linearize the exponential so that we can obtain analytical solutions of SCCs. This is applicable at asymptotically high spike-rates [42, 43] and applies well to P-type afferent spike trains because of their high baseline firing rates (about 250-300 spikes/s, [1,50,58]). To fit ISI and joint-ISI distributions of individual P-type afferents, the parameters of the dynamic threshold element *h* (*t*) (*A* and *τ*) are obtained from the afferent spike-train [42]. A single noise parameter, *a*, independent of the dynamic threshold, is obtained from the observed SCCs. These model elements and procedures allow us to determine the shaping of ISI correlations. The major results are

1. ISI correlations are determined by the auto-correlation function, **R**, of the noise process (Eqs. 12-13).
2. Non-bursting units and bursting units are described by the same functional relationship between ISI SCCs and **R** (Eqs. 12-13).
3. Non-bursting spike trains (with unimodal ISI distribution) are generated by slow noise with a decaying (positive) correlation function **R**, which in its simplest form is given by Eq. 18 (e.g., Fig. 6).
4. Bursting spike trains (with bi-modal ISI distribution) are generated by fast noise with an oscillating correlation function **R**, which in its simplest form is given by Eq. 22 (e.g., Figs. 7 and 8).
5. The two types of correlation functions are described by the sign of a single parameter, the parameter *a* of a first-order autoregressive process (Eqs 21 and 17). The AR parameter is directly related to noise bandwidth, i.e., the low or high cut-off frequencies and can be uniquely determined from the correlation, *ρ*_1_, between adjacent ISIs (Eqs. 20 and 24). More robust estimates are obtained from the ratio *ρ*_2_/ρ_1_. SCCs at subsequent lags are related to a as terms in a geometric progression (Eqs. 19 and 23).
6. While more complex patterns of SCCs can be produced by other types of noise correlation functions, only Type I and Type II SCC patterns are observed in P-type afferent spike-trains. Type III SCC patterns are mentioned here because they are commonly reported in modeling studies and are discussed further below.
7. The expression for ISI SCCs is independent of the adaptive threshold parameters (*A* and *τ*), the signal (*v*), and the firing threshold (*γ*). It is dependent only on the noise correlation function, including noise power **R**_0_.
8. The model fits ISI and joint-ISI distributions.
9. For both non-bursting and bursting units (slow and fast noise, respectively) the theoretical prediction of the sum of ISI SCCs is exactly −0.5. The sum of SCCs over all afferent spike trains is close to this limit: −0.475 ± 0.04. (*N* = 52).
10. SNR (10 log_10_(*v*^2^/**R**_0_) is generally larger than 20 dB, i.e., fluctuations in threshold (noise) are small compared to the input signal. This is in keeping with the hypothesis that spike-time jitter is small in comparison with the mean ISI.

There are two components to ISI serial correlations as is apparent from Eq. 9, where the *i*^th^ ISI is given by *t_i_* – *t*_*i*–1_ = (*γ_i_* – *γ*_*i*–1_ + *A*) /*m*. The first component is due to the difference *γ_i_* – *γ*_*i*–1_ which is coupled to the next (adjacent) interval *t*_*i*+1_ – *t_i_* by the common term *γ_i_*. This term appears with opposing signs in adjacent ISIs and hence results in a negative correlation which does not extend beyond these ISIs. If the *γ_i_* are uncorrelated then it can be shown that the adjacent ISI correlation *ρ*_1_ = −0.5 and all other *ρ_k_* = 0, for *k* ≥ 2. Thus, for independent random variables, this result follows from the property of a differencing operation and it is not indicative of memory beyond adjacent ISIs. The second component of ISI correlations is due to long-range correlations **R** in the random process *γ* which extend beyond adjacent ISIs. These correlations are endogenous, possibly biophysical in origin, and could be shaped by coding requirements. The two components to ISI correlations are made clear by restating Eq. 13 for *ρ*_1_ as

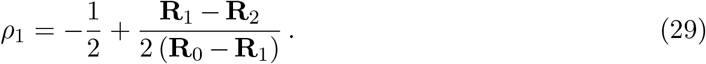

Thus, they are separable. For a wide-sense stationary process, **R**_0_ > **R**_*k*_ for all *k* ≥ 1, and so the denominator of the second term is always positive. Thus the deviation of *ρ*_1_ from −0.5 is determined by the sign of **R**_1_ – **R**_2_. This term is positive for non-bursting units (Type I SCCs), and it is negative for bursting units (Type II SCCs). The singleton case (Type III SCCs) results because noise is uncorrelated and so the term vanishes. In this case only the first component is present. For a Ornstein-Uhlenbeck or Gauss-Markov process, i.e., first order AR process with coefficient *a* > 0, **R**_1_ > **R**_2_ (Fig. 5A), and this produces a non-bursting Type I pattern. When the AR parameter is negative, i.e., the coefficient is −*a*, **R**_1_ < **R**_2_ (Fig. 5B, C), and this produces a bursting Type II pattern with a bimodal ISI distribution. Thus, a single parameter (the sign of the first-order AR parameter) can create the observed patterns of negatively correlated SCCs. Type III SCCs where *ρ*_1_ = −0.5, and *ρ_k_* = 0 for *k* ≥ 2 are trivially generated by perturbing the spike threshold with uncorrelated white noise, i.e., by setting a = 0 in the first-order AR process. We have not observed Type III neurons experimentally although some spike trains have *ρ*_1_ values close to −0.5.

The sign of the a parameter in the AR process determines the time-scale of noise fluctuations in the spike threshold, and determines the patterns of SCCs. Recall that *τ_γ_* = −*T*_1_/ln(*a*). For Type I SCCs, the AR process produces slow noise with noise bandwidth dominated by frequencies 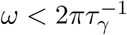 (i.e., low-pass). For Type II SCCs, the generating noise is dominated by frequencies 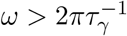 (i.e., high-pass). In the latter case, the characteristic ringing of the correlation function is distinctive, with the degree of damping being controlled by the value of *τ_γ_*.

From the above observations, we suggest that the ARMA process leading to generation of ISIs is the result of two processes: 1) An MA component which is due to a differencing operation. This contributes a value of −1/2 to *ρ*_1_, i.e., to the finite portion of the autocorrelation function (and the infinite tail of the partial autocorrelation function). 2) An AR component which is due to a feedback loop (Fig. 4). This contributes a value of −1/2 to *ϕ*_1,1_, i.e., to the finite portion of the partial autocorrelation function (and the infinite tail of the autocorrelation function). In the noise process used here, *ρ*_1_ = −1/2 ± *a*/2 (Eqs. 20 and 24 for non-bursting and bursting afferents, respectively), and so the residual contribution of the autoregressive component to *ρ*_1_ is ±*a*/2. The goodness of fit to the experimental data with a single parameter ±*a* leads us to believe that the underlying ARMA model is first-order in both the AR and MA components.

### 4.4 Comparison with other adaptation models

These results can be directly compared with results from an earlier study which modeled adaptation currents [36]. In that study, patterns of SCCs were generated by a perfect integrate-and-fire neuron under two conditions : i) a deterministic adaptation current with fast, white gaussian noise input, and ii) a stochastic adaptation current with slow, exponentially correlated (channel) noise. A deterministic adaptation current with fast noise produced patterns of SCCs that we report here as Types I and II. These patterns were characterized by a parameter *ϑ* which is analogous to the AR parameter *a* used here, with Type I pattern (positive coefficient, a) corresponding to *ϑ* > 0, and Type II pattern (negative coefficient, −*a*) corresponding to *ϑ* < 0. A pattern similar to the Type III SCC pattern reported here resulted when *ϑ* = 0. This is the same as *a* = 0 although we did not observe these neurons experimentally. While we define Type III SCCs to have only one pattern (*ρ*_1_ = −0.5, and *ρ_k_* = 0 for *k* ≥ 2) Schwalger *et al.* [36] report that *ρ_k_* = 0 when *k* ≥ 2, but the value of *ρ*_1_ is governed by an additional parameter and could take a range of values, with *ρ*_1_ = −0.5 appearing as a limiting case. The case of stochastic adaptation currents produced only positive ISI correlations which we do not observe or model here. We presume that a dynamic threshold with weak noise will not produce positive correlations for DC input, although this remains to be confirmed. In a subsequent report Schwalger and Lindner [37] extended the model to more general integrate-and-fire models and obtained a relationship between ISI SCCs and the phase-response curve (PRC). The patterns of SCCs reported here most likely correspond to their Type I PRC.

The SCCs reported here follow a geometric progression with the filter-pole a being the ratio parameter (see Eqs. 19 and 23). Schwalger *et al*. [36] and Schwalger and Lindner [37] reported that the patterns of negative correlations follow a geometric progression with the ratio parameter being *ϑ*. Similarly, a geometric progression was also reported by Urdapilleta [46]. Note that a first-order Markov process also follows a geometric progression with ratio parameter *ρ*_1_ [49, 53]. A geometric progression of SCCs is not surprising given that the ISI sequence is discrete, and thus, feedback will result in a (discrete) recurrence relation. For example, all the first-order AR processes used to predict SCCs have correlation functions which are in geometric progression (Eqs. 19, 22). Taken together with the analysis of partial correlation coefficients, and as noted above, it is likely that the underlying ARMA process generating the ISI sequence is low-order. The limiting sum of SCCs reported here (Eq. 14) is exactly −0.5, whereas [37] report a sum that asymptotically approaches a value that is slightly larger than −0.5. The average sum of SCCs in our experimental data (Fig. 2D) is also slightly larger than −0.5 (−0.475 ± 0.04.), and this merits further investigation.

In considering the results presented here using a dynamical threshold model, and those from the more general models using an adaptation current, it appears that the presence of the feedback loop (the coupling between the membrane voltage and the dynamic threshold, see Figs. 3B, 4B, and 10B) may account for almost all the properties of SCCs if the noise fluctuation is shaped appropriately. These fluctuations have the effect of reverberating around the feedback loop, creating memory and introducing negative correlations extending over multiple ISIs. However, the source of this noise is moot because it can appear in the input or the model parameters (see Eq. 31, where noise can be distributed over the input v, the threshold decay function *r*, or the spike threshold *γ*). The dynamic threshold model used here does not suggest a biophysical mechanism, but we suggest that a noisy threshold is due to an endogenous source of noise which perturbs the spiking threshold *γ* or the time-constant *τ*, i.e., noise is not passed through the input. This is an assumption, and input noise will of course shape SCCs. However, several biophysical mechanisms can account for endogenous sources of noise, including probabilistic transitions between conformational states in voltage-gated ion channels [57,59–62] leading to perturbations in gating characteristics [27,63]. These can introduce cumulative refractoriness in firing. It should be noted that the amount of noise to be added to the threshold is small, generally weaker than 20 dB SNR (see Figs. 6, 7, and 8). That only weak noise may be necessary has been reported earlier in [37], and in a study on threshold shifts [57].

While a dynamic threshold model does not explicitly incorporate biophysics, models with adaptation currents can be more readily tied to biophysical conductances. An ion channel that may be a substantial contributor to adaptation currents is the non-inactivating M-current (KCNQ/Kv7 family) [64] which is a voltage-gated potassium channel with slow dynamics (relative to the mean ISI). This channel may be responsible for spike-timing precision, and hence a timing-based neural code (see [42] for a discussion). The possible role of the M-current and the specific sources of noise have been explored earlier (see [36] above in relation to types of SCCs, and [5]). Both studies [5, 36] included a modified conductance-based Traub-Miles model [65, 66] frequently used in modeling M-currents. They concluded that negative correlations were a consequence of a deterministic adaptation current with fast white Gaussian noise input rather than a stochastic adaptation current with slow correlated noise. The current report shows that at least some of the results with adaptation currents can be reproduced with a simple dynamic threshold with a noisy spiking threshold or a noisy decay time-constant. Further, either slow noise or fast noise can produce negative correlations if injected into the spiking threshold, with the time-scale of noise fluctuations determining the type of SCC pattern that is produced (i.e., bursting or non-bursting). In another study [27] the first SCC (*ρ*_1_) was compared when the generating models were a leaky integrate-and-fire (LIF) neuron with an adaptation current (LIFAC) or a dynamic threshold (LIFDT). When the LIFAC and LIFDT models are matched so that the spike-rate and adaptation are about the same, the ρ1 as a function of spike rate are also similar. This gives reason to believe that either models of adaptation can give rise to similar patterns of SCCs, possibly due to the simple feedback present in the models. However, a more detailed biophysical investigation is needed before we can tease apart the differences at a mechanistic level.

### 4.5 Power spectra and noise-shaping

Negatively correlated spike trains have implications for information transmission [23]. These spike trains have a power spectra *P* (*ω*) that rolls off towards DC [24,37], effectively reducing low-frequency noise and improving SNR in the baseband (also referred to as noise-shaping). The connection between DC power *P* (*ω* = 0) and the ISI SCCs is given by Eq. 16 [49], where it can be seen that negative correlations have a tendency to reduce noise at zero-frequency. From the theoretical limit of the sum of SCCs (−0.5) reported here in Eq. 14, similar to the results with adaptation currents [37], DC-power vanishes, thus yielding a perfect DC block allowing low-frequency signals to be transmitted with very high SNR. The limiting sum of observed SCCs is only just a little larger than the optimum sum (−0.5), and this allows some noise to bleed through at zero-frequency. These results are in line with the results on power spectra first predicted by Chacron *et al.* [23,24] and more recently extended to models with adaptation currents [37].

### 4.6 Implications for optimal coding

The model presented here sets the spike threshold *γ* = *A*/2. We showed previously that the time-varying dynamic threshold *r* (*t*) is an estimator of the input signal *v*(*t*), and the mean-squared estimation error ((*v*(*t*) – *r*(*t*))^2^) is minimized when the firing threshold is *γ* = *A*/2 [42,43]. The error increases for any other value of *γ*, in particular *γ* = 0 which is the value usually adopted in the literature. Thus, for optimal timing of spikes the ongoing estimation error must be bounded below by *γ* = *A*/2. When this bound is reached, the neuron fires a spike and raises the adaptive threshold variable by *A*. Viewed in this manner, the spike generator directly encodes the error and not the signal (Figs. 3B, 4B). This is much more efficient than directly encoding the signal (or some filtered version of the signal). The coding error is equivalent to quantization error in digital coding. This coding mechanism is analogous to one-bit delta modulation [67], but it is asymmetric because the threshold is one-sided. Thus, coding with an dynamic threshold is analogous to lossy digital coding or source coding. That is, a neuron performs data compression.

For a constant DC-valued input, the SCCs generated by the model are dependent solely on noise correlations **R**, and are independent of all other parameters, including the dynamic threshold parameters *A* and *τ*, the mean value of the threshold *γ*, and the constant DC-valued input signal *v*. The model parameters *A* and *τ* are fitted from the baseline (spontaneous) spike-train through the relationship *Aτ* = *vT*_1_, where *v* is the bias input and *T*_1_ is the long-term mean ISI. This allows us to reduce the degrees of freedom to one, to either *A* or *τ*, and then determine the free parameter by optimizing the model spike train to match experimental spike-times (see [42] for details of the procedure). Thus, the dynamic threshold parameters are not estimated from the observed SCCs. This fits with our understanding of the dynamic threshold as an optimal estimator with its parameters being determined by coding quality and an energy constraint [42, 43]. When the deterministic parameters are fixed, a noisy threshold can be generated by estimating the filter parameter a from *ρ*_1_ or *ρ*_2_/*ρ*_1_ to match the observed SCCs. This decoupling of dynamic threshold parameters from SCCs implies that a correlation function **R** can be generated from a specified sequence of SCCs (determined, say experimentally) without reference to the time-varying threshold. Thus, a family of spike trains with arbitrary mean ISI, ISI and joint-ISI distributions can be generated, all of which have the same sequence or pattern of SCCs. Some iteration will be needed to fit the ISI and joint-ISI distributions from the dynamic threshold parameters A and *τ* as is done here, but this is not too difficult.

### 4.7 Further work and conclusion

The main goal of this work was to show that a simple dynamic threshold model with few parameters can reproduce known negative correlations, and further, model experimental data and provide insight into the origin and patterns of ISI negative correlations. We proceeded on the assumption that there is only one experimentally observable quantity: the sequence of spike times. Based on spike-times, we have shown that the ISI distribution, the joint-ISI distribution, and SCCs can be predicted accurately. The approach used here suffers from several drawbacks. The mean firing rate, or equivalently mean ISI, is implicitly defined through the model parameters and hence it does not influence SCCs (the noise correlation function is discretized using ISI lag-number, and not the absolute value of the ISI). The SCCs obtained with this approach are thus invariant under firing rate. In reality, we expect correlations to be small when the spike rate is high or low, reaching a maximum at some intermediate firing rate (see [27]). The independence of SCCs from mean ISI is a result of infinite memory in the model because the exponential threshold was approximated as a line with slope m. A second drawback is that the input is a fixed DC-value and does not incorporate dynamics or noise. It should be noted that noise in the current model is assumed to perturb the spiking threshold, however, it can be exogenous, i.e., present in the input *v*, or present jointly in *γ* and *v*. Currently, there is no way to distinguish between these cases (see the deterministic firing rule specified in Eq. 7) because noise can be distributed over *γ* or over *v*. Thus, for a noisy DC-valued input and fixed *γ*, the results will be the same as for a noisy *γ*. Rather than determining the effect of input dynamics on SCCs an alternative and promising line of enquiry is to determine the role of ISI SCCs, (determined under spontaneously active conditions) on processing time-varying inputs of varying bandwidth, i.e., on signal encoding. As noted earlier [42], the bandwidth of the input will affect the quality of the coding and the extent to which ISI correlations maintain coding fidelity (i.e., the stimulus estimation error). To determine these effects, the model can be readily adapted to non-constant inputs by making the input piece-wise linear between spikes. This approach and the necessary theory was developed earlier with a deterministic version of the dynamic threshold model to predict spike-times [42–44]. We hypothesize that a stochastic extension of the model with weak noise (added to the spike threshold or decay time-constant) should not disrupt timing other than introduce timing jitter. This has been shown in neocortical neurons where spike-timing is reliable across repeated presentations of rapidly fluctuating time-varying signals [68], and can be accurately modeled using a dynamic threshold [42]. Another study, albeit preliminary, showed that the patterns of negative correlations do not change when there is a time varying input [69]. While it is known that negative correlations, and more generally adaptation currents can reduce variability and stabilize firing rate in single neurons (see for example, [1,37]) and populations of neurons [21,70], robust spike-times (i.e, reduced timing jitter) in response to time-varying signals may be yet another benefit provided by negative SCCs in ISIs. Thus, the use of a more exact, i.e., leaky, dynamic threshold function, the extension to time-varying inputs, and the determination of the linkages between dynamics of the input, the threshold, and coding fidelity, are the obvious next steps.

The simplified dynamic threshold model used here demonstrates the influence of noise on ISI correlations. We provide a closed-form expression for SCCs as a function of the noise correlation **R**. This is useful for solving the inverse problem when SCCs are known and the discrete sequence **R**_*k*_ is to be determined. The forward problem is also readily solved, i.e., given a sequence **R**_*k*_, the SCCs can be evaluated. We illustrate with a first-order AR model, and show that a single-parameter (the sign of the coefficient in the first-order AR process) captures the pattern of observed SCCs, and in this respect it agrees with a detailed dynamical model using adaptation currents [37]. This model can fit observed spike-times with good accuracy [40, 42], is energy-efficient while maximizing coding fidelity [42, 43], and is computationally inexpensive. Finally, the model provides a quick and efficient way to generate surrogate data mimicking negatively correlated spike trains observed in experimental neurons. Thus, the model can be useful for understanding timing-based neural codes.

## Appendix

### Deterministic and stochastic adaptive threshold model

In the deterministic adaptive threshold model (Fig. 3A), if *t*_*i*–1_ and *t_i_*, *i* ≥ 1, are successive spike-times, then the time evolution of the adaptive threshold *r* (*t*) is given by

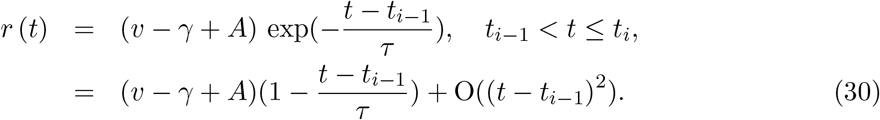

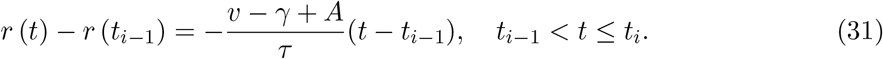

Let *m* = (*v* – *γ* + *A*) /*τ* > 0, so that the slope of the adapting threshold is −*m*, then

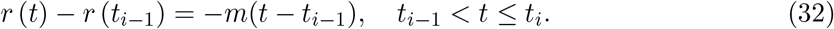

Noting that *r*(*t_i_*) = *v* – *γ*,

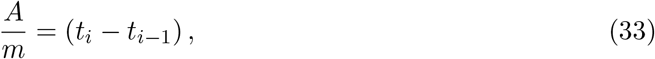

which is the deterministic firing threshold for a constant, DC-level signal.

### Dynamic threshold with random firing threshold

We will now provide a stochastic extension to the model by making the spike threshold (*γ*) noisy while leaving all else unchanged (Fig. 4). Let *γ* be a discrete wide-sense stationary process with mean **E**[*γ*], discrete auto-correlation function **R**_*k*_ and power **E** *γ*^2^ = **R**_0_. The spike threshold with additive noise assumes the value *γ_i_*, *i* ≥ 1 immediately after the (*i* – 1)^th^ spike and remains constant in the time interval *t*_*i*–1_ < *t* ≤ *t_i_* (Fig 4A) [24,48]. Thus, the *i*^th^ spike is emitted when the error satisfies the condition

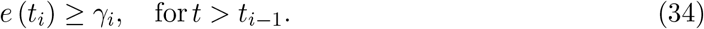

Subsequently the adaptive threshold jumps to a higher value specified by *v* – *γ_i_* + *A*, and the noisy spike threshold assumes a new value *γ*_*i*+1_. From Fig. 4A, proceeding as before

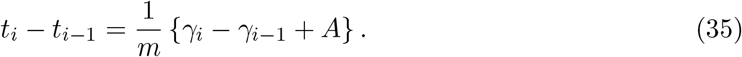

The mean ISI is therefore **E** [*t_i_* – *t*_*i*–1_] = *A/m* as in the deterministic case given by Eq. 7. Thus the correlation function for the ISI sequence is

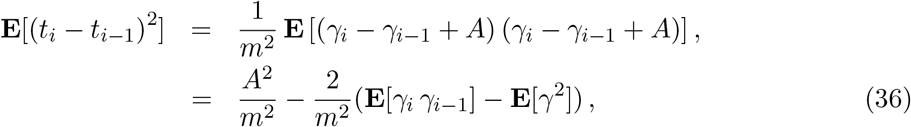

with covariance

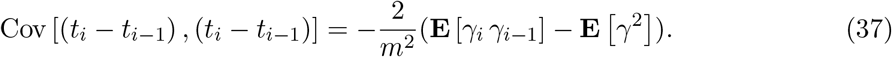

More generally, for the sum of *k* ISIs

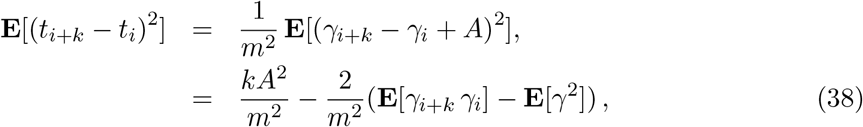

with covariance

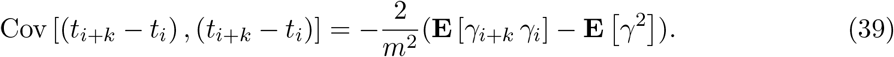

From Eqs. 36 and 38 it can be seen that there are two terms in each of these ISI correlation functions. The first is due to the input signal (which gives a mean ISI of *A/m*) and the second is due to the discrete-event noise process *γ*. From the assumption of wide-sense stationarity the auto-correlation function **E** [*γ_i_ γ_j_*] can be written as **R** (*t_i_* – *t_j_*). The discrete nature of the correlation function **R** is made clear in the following way. Denote the mean of the *k*^th^-order interval by *T_k_* = **E**[*t*_*i*+*k*_ – *t_i_*], then the mean ISI is *T*_1_ (= *A/m*), and further *T_k_* = *kT*_1_. The random variable *γ* is generated once every ISI and thus, the discrete auto-correlation function **R** takes values at successive multiples of the mean ISI, i.e., **R** (*T*_1_), **R** (*T*_2_), etc., and will be denoted by **R**_1_, **R**_2_, etc., respectively. As noted before, **R**_0_ is noise power. That is, we can write

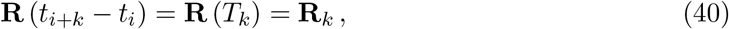

and therefore

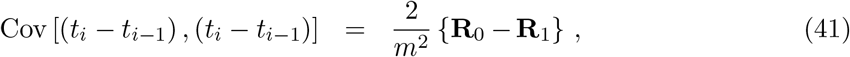

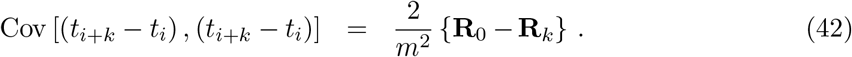

The cross-covariance of two ISIs (*t_i_* – *t*_*i*–1_) and (*t*_*i*+*k*_ – *t*_*i*+*k*–1_) separated by lag *k* ≥ 1 is

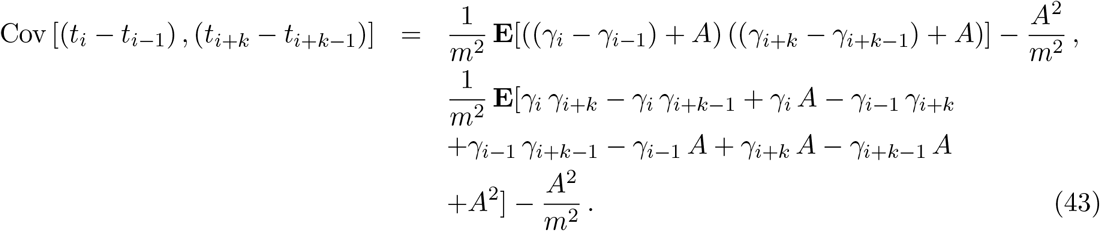

This yields

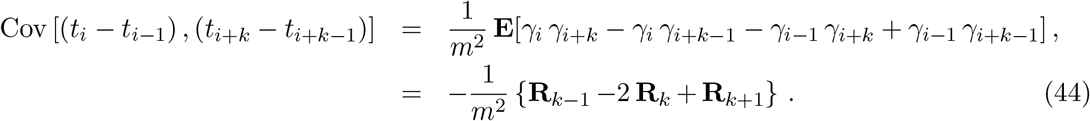

The general formula for serial correlation coefficients at lag *k* is [49],

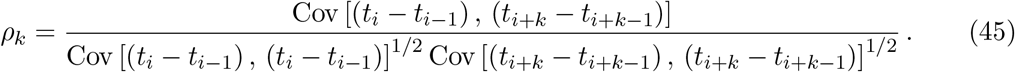

For a wide-sense stationary process the covariances are constant, and the subscript *i* can be dropped. Introducing Eqs. 41 and 44 into Eq. 45 yields

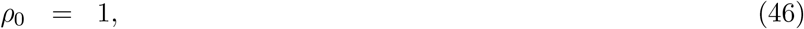

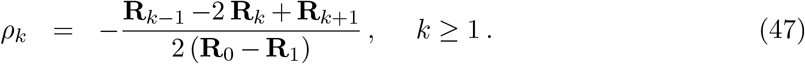

These equations are reproduced in Results as Eqs. 12 and 13, respectively. The serial-correlation coefficients (SCCs) given by Eqs. 46 and 47 are independent of the slope m of the reconstruction filter (the decay rate of the adaptive threshold) and the reconstruction filter gain A. Thus, the observed correlation structure of the spike-train is determined solely by the noise statistics of the firing threshold *γ*.

### Dynamic threshold with a random time-constant

We begin with Eqs. 9, 25, and 26, which transform a random firing threshold into a random time-constant *τ_i_* that is held constant between spike-times [*t*_*i*–1_, *t_i_*) with mean *τ* = **E** [*τ_i_*] (Fig. 10). It is immediately apparent that the covariance of the sequence *τ_i_* (sampled at the ISIs) is the same as the covariance of the ISI sequence up to a scale-factor, and therefore the serial correlations of *τ* (*ρ_τ,k_*, *k* ≥ 0) are the same as the serial correlations of the ISIs. We can establish this as follows. From Eq. 26

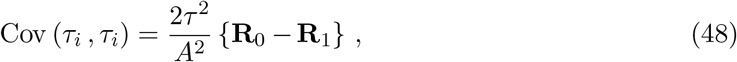

and

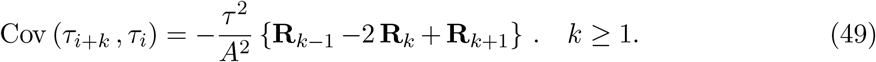

From Eqs. 45, 48 and 49 we can determine the serial correlation coefficients of the random time-constants to be

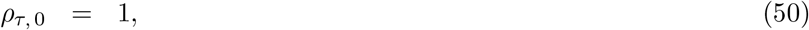

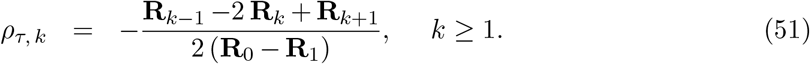

These equations are reproduced in Results as Eqs. 27 and 28, respectively. The right side of the above equations, Eqs. 50 and 51, are the same as Eqs. 46 and 47, which are the expressions for the SCCs of a spike train generated with a noisy adaptive threshold. Thus, the random filter time-constants have the same serial correlation coefficients as the interspike intervals, *ρ_τ,k_* = *ρ_k_*. This is a “pass through” effect where the correlations observed in the time-constant are directly reflected in the ISI correlations.

In summary, a noisy threshold or a noisy dynamic threshold time-constant can generate spike trains with the same ISIs and SCCs, and either method can be used.

### Sum of serial correlation coefficients

We now determine the limiting sum of the SCCs over all lags. Let us first transform *ρ_k_* as follows by defining

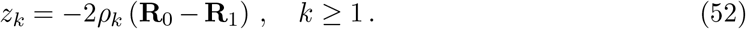

Then from Eq. 47, *z_k_* is a moving-average process such that

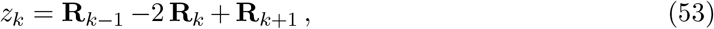

from which it is easy to show that

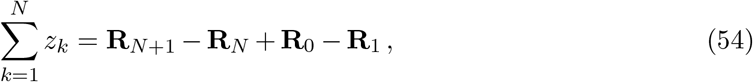

where *N* is the *N*^th^-order interval. We make the assumption that the process *γ* is aperiodic and the auto-correlation function **R** (*N*) → 0 when *N* → ∞, and so (**R**_*N*+1_ – **R**_N_) → 0. That is, the process decorrelates over long time-scales. Thus,

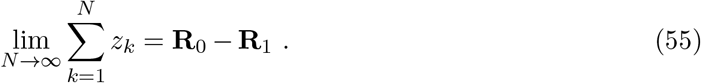

By summing Eq. 52 over all *k*, and inserting the result from Eq. 55, the limiting sum of the ISI serial correlation coefficients for the spike-train is

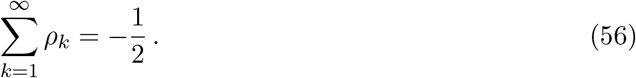

## Acknowledgments

The authors were partially supported by National Science Foundation grants EFRI-BSBA-0938007 and IGERT 0903622. Additional funds and support were provided by the College of Engineering and Coordinated Science Laboratory (UIUC), the Advanced Digital Sciences Center (Illinois at Singapore), and Ahmedabad University. Electric fish data were collected in the laboratory of Mark E. Nelson, UIUC, through a National Institutes of Health grant R01MH49242 and National Science Foundation grant IBN-0078206.

## Notes

### Competing Interest Statement

The authors have declared no competing interest.

### Summary of Updates

This version of the manuscript addresses reviews questions, comments, and suggestions.

